# Preventing peritendinous adhesions using lubricious supramolecular hydrogels

**DOI:** 10.1101/2025.02.22.639655

**Authors:** Emily L. Meany, Christian M. Williams, Ye Eun Song, Vanessa M. Doulames, Sophia J. Bailey, Shoshana C. Williams, Carolyn K. Jons, Paige M. Fox, Eric A. Appel

**Affiliations:** Department of Bioengineering, Stanford University, Stanford, CA, 94305, USA; Department of Material Science & Engineering, Stanford University, Stanford, CA, 94305, USA; Sarafan ChEM-H, Stanford University, Stanford, CA 94305, USA; Department of Chemistry, Stanford University, Stanford, CA, 94305, USA; Robert A. Chase Hand and Upper Limb Center, Stanford University Medical Center, Stanford, CA, 94305, USA; Division of Plastic and Reconstructive Surgery, Stanford University School of Medicine, Stanford, CA, 94305, USA; Division of Plastic Surgery, Veterans Affairs Palo Alto Health Care System, Palo Alto, CA, 94304 USA; Wood Institute for the Environment, Stanford University, Stanford CA 94305, USA; Department of Pediatrics (Endocrinology), Stanford University, Stanford CA 94305, USA

**Author notes:** These authors contributed equally to this work (Emily L. Meany, Christian M. Williams).

**Keywords:** Adhesions, Hydrogel, Flexor Tendon, Surgery, Adhesion Barrier

## Abstract

Of the 1.5 million emergency room visits each year in the United States due to flexor tendon injuries in the hand, over 30-40% result in peritendinous adhesions which can limit range of motion (ROM) and severely impact an individual’s quality of life. Adhesions are fibrous scar-like tissues which can form between adjacent tissues in the body in response to injury, inflammation, or during normal healing following surgery. Currently, there is no widespread solution for adhesion prevention in the delicate space of the digit while allowing a patient full ROM quickly after surgery. There is a clear clinical need for a material capable of limiting adhesion formation which is simple to apply, does not impair healing, remains at the application site during motion and initial inflammation (days - weeks), and leaves tendon glide unencumbered. In this work, we developed dynamically crosslinked, bioresorbable supramolecular hydrogels as easy-to-apply lubricious barriers to prevent the formation of peritendinous adhesions. These hydrogels exhibit excellent long-term stability, injectability, and thermally stable viscoelastic properties that allow for simple storage and facile application. We evaluated interactions at the interface of the hydrogels and relevant tissues, including human tendon and skin, in shear and extensional stress modes and demonstrated a unique mechanism of adhesion prevention based on maintenance of a lubricious hydrogel barrier between tissues. *Ex vivo* studies show that the hydrogels did not impair the gliding behavior nor mechanical properties of tendons when applied in cadaveric human hands following clinically relevant flexor tendon repair. We further applied these hydrogels in a preclinical rat Achilles tendon injury model and observed prolonged local retention at the repair site as well as improved recovery of key functional metrics, including ROM and maximal dorsiflexion. Further, these hydrogels were safe and did not impair tendon strength nor healing compared to the current standard of care. These dynamic, biocompatible hydrogels present a novel solution to the significant problem of peritendinous adhesions with clear translational potential.

## 1. Introduction

Post-operative adhesions represent a major challenge to rehabilitation following surgery. Adhesions are fibrous bands of scar-like tissue, ranging in severity from wispy filaments to thick and vascularized connections, which attach previously separate tissues and form as a consequence of injury and inflammation during normal wound healing following surgical interventions. Adhesions are important contributors to complications after a wide range of surgeries, from intra-abdominal surgery where they can cause small bowel obstructions and increased re-operation times, to orthopedic surgery where they often limit functional recovery.^1-6^ Peritendinous adhesions are particularly insidious as these directly restrict normal range of motion (ROM) after tendon repair, can cause severe pain, and impair patient quality of life.^1,5-8^ In the United States alone there are 1.5 million emergency room visits per year due to flexor tendon injuries and more than 30 million people worldwide suffer from tendon injuries every year, accounting for over $140 billion in healthcare costs.^9^ Of these injuries, more than 30-40% result in peritendinous adhesions which reduce ROM and grip strength, and may be severe enough to require additional surgery to remove the adhesions.^1^

The only available treatment for peritendinous adhesions after they are formed is tenolysis, a follow-up procedure wherein adhesion tissue is surgically removed. Unfortunately, this procedure can maximally restore ROM to 80% of pre-injury ROM and risks inducing more adhesions.^10-12^ Considering the prevalence, risks, and lack of non-invasive treatments for peritendinous adhesions, a prophylactic intervention would be ideal. However, there are currently no well-studied, widely used products for the prevention of peritendinous adhesions in the United States.^13^ As a result, patients and providers rely on early post-operative mobilization to help limit adhesion formation after repair and tenolysis, sometimes beginning on the same day as the surgery.^14,15^ These mobilization regimens help to activate intrinsic healing of the tendon and reduce formation of thick adhesions, but they must balance increased risk of repair rupture and their efficacy relies on patient compliance, which wanes over time.^1,7,14,15^ An alternative solution, employed first for abdominal adhesions, is the use of a barrier to separate healing tissues and prevent adhesions. Such an intervention could be beneficial for many orthopedic applications, such as injuries to flexor tendons, the rotator cuff, and the Achilles tendon. The only FDA-approved products for peritendinous adhesion prevention are Tenoglide (Integra Lifesciences, NJ, USA) and Versawrap (Alafair Biosciences, TX, USA), both of which are polymer sheets that lack detailed clinical data for their effectiveness.^13,16-19^ Depending on surgical location, it can be difficult to wrap a tendon in a sheet and it is likely that sheet would dislodge during movement or otherwise prevent early mobilization. For an adhesion barrier to be efficacious, it must completely cover the injured tissue during critical inflammatory and proliferative periods of wound healing, roughly 2-3 weeks for tendons, without limiting movement nor impairing the speed and quality of healing.^1,5-7,20^

To attain these results, dynamic hydrogels have been investigated as anti-adhesion barriers. An ideal hydrogel for preventing peritendinous adhesions would be (i) shear-thinning to be easily spread for complete tissue coverage and readily incorporated into existing surgical procedures, (ii) tissue adherent to ensure local retention after application, (iii) viscoelastic to not hinder the motion of dynamic tissues, and (iv) biocompatible to not impair healing. Our team has previously published a novel approach to prevent post-operative adhesions leveraging a dynamic, supramolecular polymer-nanoparticle (PNP) hydrogel which exploits non-specific, multivalent interactions between cellulose-derived polymers and polymeric nanoparticles.^21^ Prior studies showed a significant reduction in post-operative adhesions in both the pericardial and peritoneal spaces after use of a PNP hydrogel barrier, demonstrating efficacy of dynamic hydrogels as anti-adhesion barriers.^22,23^ Based on this work, we sought to design a hydrogel suited for peritendinous applications, where a great unmet clinical need remains.

To this end, our lab has developed a novel hydrogel platform composed of a hydrophobically modified cellulose derived polymer (HPMC-C_18_) and a surfactant excipient commonly used in pharmaceutical products Tween 20 (Tw20).^24^ Major advantages of these polymer-tween (PTw) formulations are their ease of synthesis and scalability. Whereas PNP hydrogel manufacturing involves several steps, including synthesizing each polymeric component, preparing the self-assembled nanoparticles, and finally mixing the hydrogel, PTw hydrogels can instead be generated from commercially available materials with one mixing step. The PTw hydrogel platform allows for facile tuning of stiffness and viscoelasticity by adjusting concentrations of HPMC-C_18_ and Tw20. These hydrogels exhibit similar mechanical behaviors to PNP hydrogels, including shear thinning, facile injectability at the site of repair, and robust tissue adherence, allowing them to form lubricious barriers between tendon and other tissues while simultaneously allowing free tendon gliding. In this work, we optimized PTw hydrogels for use in prevention of peritendinous adhesions and used shear and extensional rheology to characterize their yielding properties on relevant murine and human tissues. We show feasibility of PTw hydrogel use in human cadaver hands following clinically relevant flexor tendon injury and repair, demonstrating the material is easy for a surgeon to apply following surgery and does not impede gliding nor otherwise reduce tissue integrity. We then evaluated the efficacy and safety of these materials in a preclinical rodent Achilles tendon injury model and found improved functional recovery at two months as well as no negative impact on tendon healing.

## 2. Results

### 2.1 Characterization and stability of dynamic hydrogels

The current standard of care following flexor tendon injury is immediate primary repair of the tendon, followed by a regimen of early mobilization to promote intrinsic and limit extrinsic healing to reduce peritendinous adhesion formation. Yet, early mobilization timing and regimen are debated among surgeons and adhesion formation remains a major concern, particularly in the frequently injured zone II of the hand where extrinsic healing is supported by the tendon synovial sheath.^1,25,26^ Once formed, adhesions between the tendon and surrounding tissues limit tendon movement and digit range of motion. In contrast, we propose a care solution wherein the surgeon applies a dynamic hydrogel around the tendon following repair, before skin closure. This material will form a lubricious barrier separating the tendon from surrounding tissues, limiting adhesion formation, while simultaneously allowing free tendon glide (**Figure 1A**).

**Figure 1.**
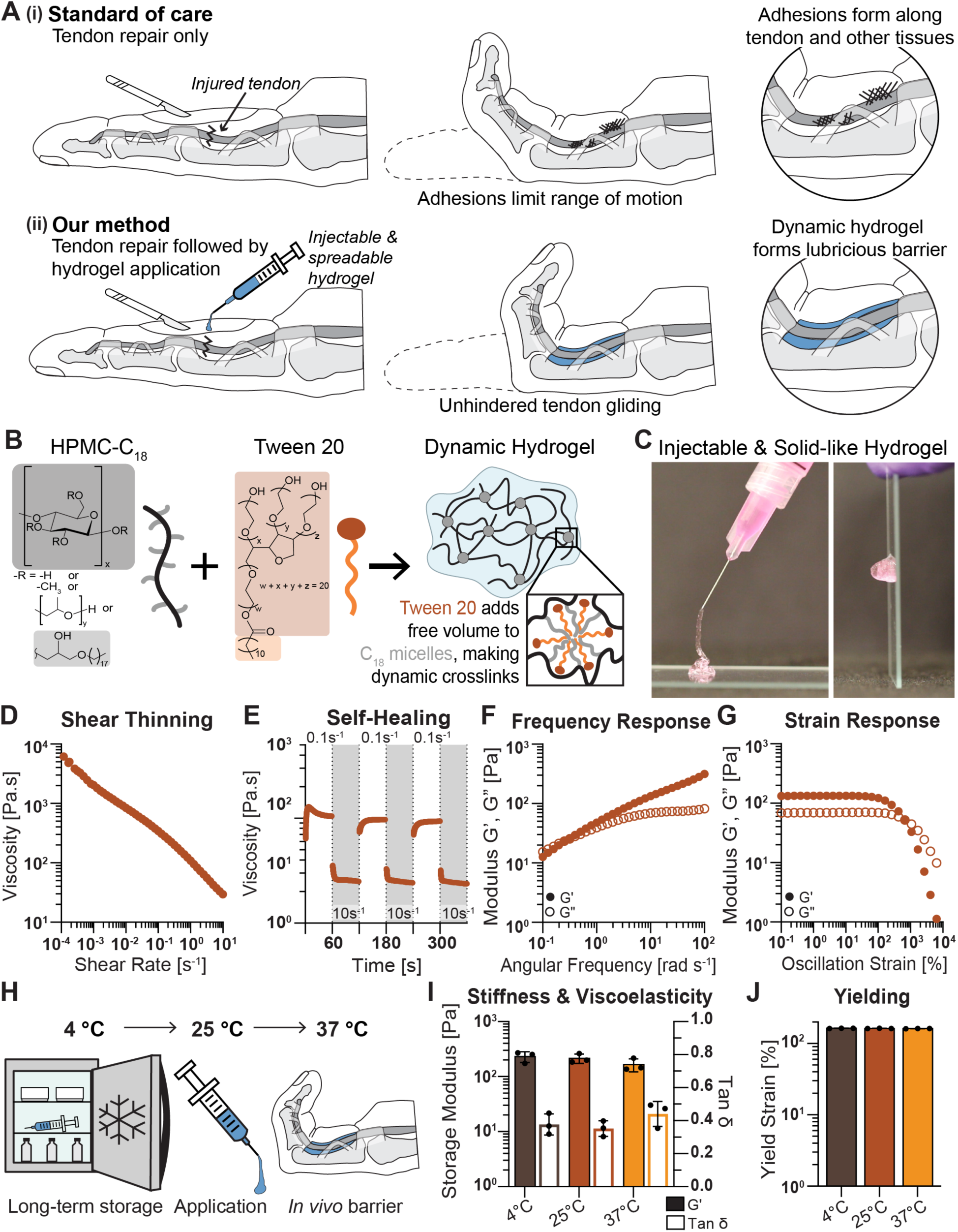
Characterization of PTw hydrogel. A) Schematic of current standard of care treatment for injured flexor tendons in the hand compared with our proposed treatment. PTw hydrogel applied at the site of repair before surgical closure forms a lubricious barrier, allowing unhindered tendon gliding during healing and limiting tissue-tissue adhesion formation. B) Chemical composition of dynamic PTw hydrogel. Tween 20 surfactant introduces free volume into HPMC-C_18_ crosslinking nodes, forming a dynamic hydrogel. C) PTw hydrogel is injectable through small gauge (30 gauge) blunt-tipped needles and recovers as a solid-like depot following injection. D) High-to-low shear rheology shows PTw hydrogels are shear-thinning, and E) step-shear rheology shows the viscosity drops an order of magnitude under high shear (10 s^-1^) and recovers rapidly at low shear (1 s^-1^) over repeated cycles. F) Frequency sweep shows PTw is dynamic and solid-like (G′ larger than G″) over relevant time scales. G) Amplitude sweep shows PTw is solid-like over relevant strains. H) Schematic of temperatures PTw hydrogels are subject to through storage and application. I) G′ and tan δ at 40 rad·s^-1^ and J) yield strain (strain where G′ < 0.85 * max(G′)) for PTw at 4, 25, 37 °C. Data shown as mean ± SD, n=3.

We developed a dynamic hydrogel system composed of stearyl-modified hydroxypropyl methylcellulose polymers (HPMC-C_18_) and tween 20 surfactant (Tw20) (**Figure 1B**).^24^ When mixed, Tw20 incorporates into existing C_18_ sidechain interactions among the HPMC-C_18_ polymers, adding free volume and producing more dynamic crosslinks.^24^ The resulting PTw hydrogels were easily injected through blunt-needles as small as 30-gauge and reformed as robust solid-like materials following injection (**Figure 1C**). In this work, we engineered a PTw hydrogel formulation comprising 1.8 wt% HPMC-C_18_ and 0.5% Tw20 to closely match important rheological properties of a previously developed PNP-1-10 hydrogel formulation (1 wt% HPMC-C_12_ and 10 wt% PEG-PLA nanoparticles; **Figure S1**) that was found to be beneficial in limiting adhesions in the cardiac and peritoneal spaces.^22,23^

Rheological characterization of these PTw hydrogels showed robust shear-thinning and self-healing behavior, with viscosity dramatically reduced by over an order of magnitude at high shear rates, followed by quick recovery during repeated cycling between low and high shear rates (1 and 10 rad s^-1^) (**Figure 1D, E**). These properties indicate PTw hydrogels can withstand high-shear events like injection or digit flexion and recover their pre-shear mechanical properties repeatedly. PTw hydrogels exhibit solid-like mechanical properties in frequency sweep rheological experiments, with G′ storage modulus larger than G″ loss modulus over several decades of frequency space (**Figure 1F**). These materials also exhibit an exceptionally broad linear viscoelastic region (i.e., mechanical properties are independent of strain) up to almost 200% strains, after which point the materials begin to yield and flow (**Figure 1G**). For reference, the shear strains experienced in the tendon sheath can exceed 1000%.^27-29^ We hypothesized the shear-thinning behaviors under these conditions would reduce viscous drag during movement of the digit and the rapid self-healing behavior would allow the hydrogels to be retained at the application site even after large strains are applied.

For use in peritendinous adhesion prevention, it is important for PTw hydrogels to maintain key mechanical properties at the different temperatures the materials will be exposed to, from 4 °C refrigeration during storage to 25 °C during application and 37 °C when in the body (**Figure 1H**). We conducted frequency and strain amplitude sweeps at each of these temperatures, extracted the storage modulus, tan δ, and yield strain values, and found PTw mechanical properties were invariant to temperature in this range (**Figure 1I, J**). We further stored materials at 4 °C for over a year and tested them at 25 °C and found these properties were unchanged, indicating PTw hydrogels can be stored for prolonged periods without degradation (**Figure S2**).

### 2.2 Tissue adhesive properties of PTw hydrogels

One important property of an adhesion barrier is its ability to adhere to tissue and be retained at the surgical site after application, maintaining a lubricious barrier between tissues. This property is particularly relevant in the peritendinous space, where a hydrogel would regularly experience strains as digits flex and extend. Under shear stress, cohesive yielding of the hydrogel is desirable and indicates good adhesion at the hydrogel-tissue interface, whereas adhesive failure at the interface is undesirable and would lead to hydrogel dislodging from the application site over time by slipping against tissue motion (**Figure 2A**). To probe this behavior, we assessed the yield stress of PTw hydrogels on a standard serrated parallel plate geometry and three relevant tissues: human hypodermis, human tendon, and murine hypodermis (**Figure 2B**). In the event of adhesive failure on tissue, slippage would cause the apparent yield stress to be less than the value measured on a standard geometry. We found the yield stress for PTw hydrogels on all tissue substrates was greater than or equal to that on a standard geometry, indicating cohesive yielding under shear deformation (**Figure 2E-F**). Additionally, we investigated the tissue adhesiveness of PTw hydrogels under normal stress with filament stretching extensional rheometry. In the case of cohesive yielding, the hydrogel filament should break at a given strain, but in the undesirable case of adhesive failure, the hydrogel will separate from the substrate (**Figure 2C**). We conducted these experiments on a standard geometry, as well as human hypodermis, human tendon, and murine hypodermis, and observed cohesive yielding on all substrates, highlighting that PTw hydrogel will adhere to the tissue it is applied to even if it is pulled away from it (**Figure 2D**). Further, the extensional strain at break was roughly equivalent for all tested surfaces, with PTw hydrogel extending to upwards of 16 times its original length (**Figure 2G**). We also tested the tissue adhesive properties of our previously-reported PNP hydrogels and confirmed they exhibit similar behaviors to PTw hydrogels, likely due to the broadly tissue adherent properties of the HPMC polymers comprising both of these gels (**Figure S3**).^30,31^ The demonstrated tissue adhesive properties of PTw hydrogels ultimately promote their local retention and performance as lubricious anti-adhesion barriers.

**Figure 2.**
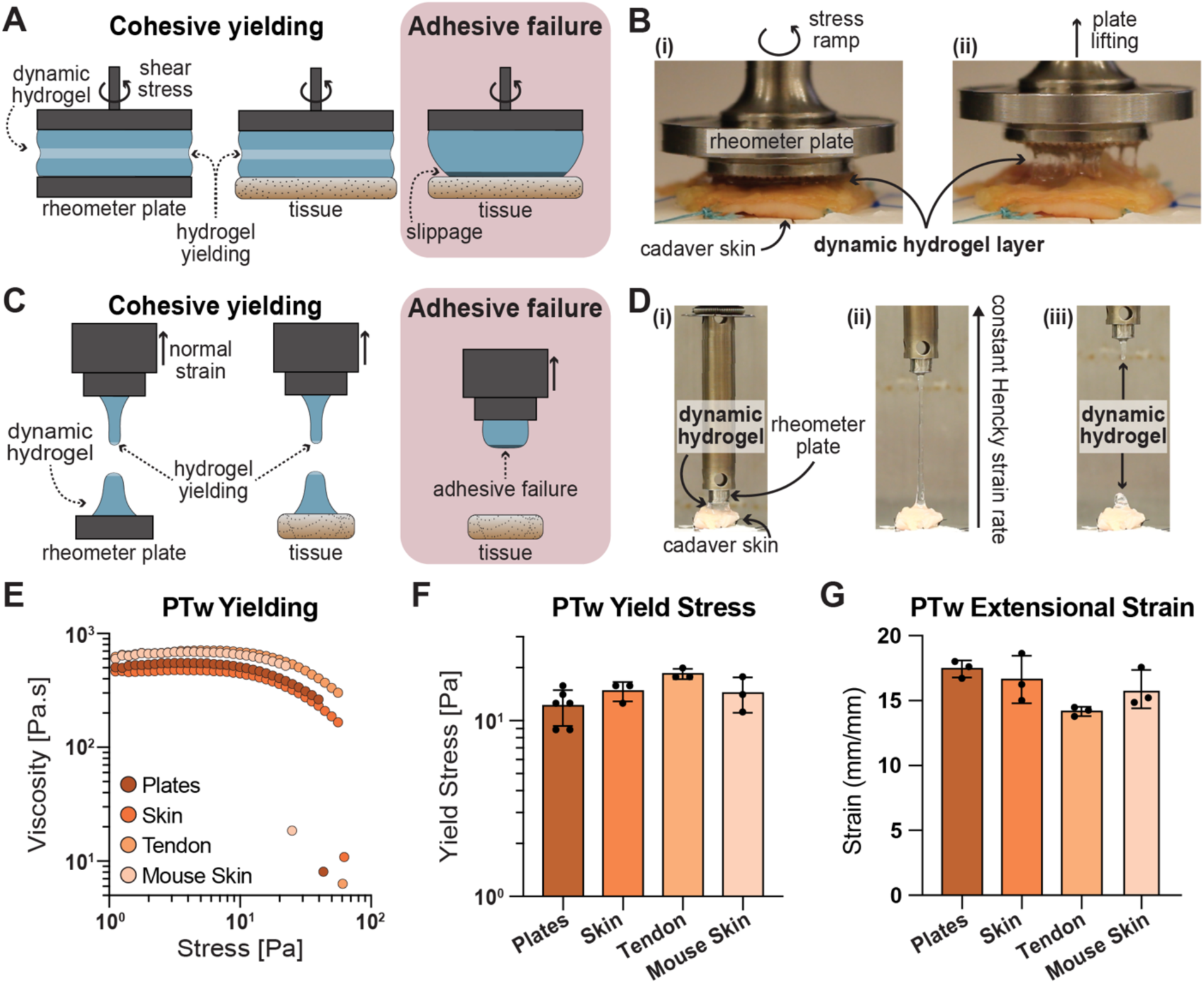
Tissue adhesiveness of PTw hydrogel. A) Schematic demonstrating cohesive yielding vs. adhesive failure of PTw hydrogel under shear stress. B) Photos showing (i) shear stress ramp on human cadaver tissue and (ii) lifting geometry after stress ramp experiment. C) Schematic demonstrating cohesive yielding vs. adhesive failure under extensional stress. D) Photos of (i) extensional rheometry of PTw hydrogel during (ii) extension and (iii) cohesive yielding of PTw. E) Yielding behavior of PTw hydrogel in a serrated parallel plate geometry and on various tissue substrates in a stress-ramp experiment with F) quantified yield stress values (defined at a 15% decrease from the peak viscosity). G) Extensional strain at break for PTw hydrogels before cohesive failure with various substrates. Data shown as mean ± SD, n=3.

### 2.3 PTw hydrogel compatibility in human cadaver digits

A key concern when introducing material to limit peritendinous adhesions is that the space surrounding tendons in the digit is limited by the tendon sheath and any physical barrier should be able to coat a tendon while allowing it to glide freely within the pulleys. To evaluate these behaviors, we performed clinically relevant zone II flexor tendon repair in human cadaver arms and evaluated several metrics to determine how applied PTw material impacted tendon glide. The flexor digitorum profundus (FDP) tendon was visualized and transected between the annular pulleys A2 and A4, then repaired with a modified Kessler stitch and a horizontal mattress to recapitulate the most common clinically observed zone II flexor tendon injuries with a 4-strand repair (**Figure 3A**).^32,33^ For PTw treated digits, hydrogel was injected at the repair site and spread to ensure full coverage of the FDP tendon and repair on all sides, including underneath at the interface with the flexor digitorum superficialis (FDS) tendon, then the skin was sutured closed. For better visualization in these experiments, PTw hydrogels were formulated with rhodamine-tagged HPMC-C_18_, giving the PTw hydrogel a pink color that was easily distinguishable from the surrounding tissue. Cadaver arms were set in a custom rig which connected the FDP tendon to a material testing system (MTS) on one side and the target digit to a counterweight at the other (fingertip) (**Figure 3B**). The MTS measured the encountered forces as it moved up, flexing the digit, and down, extending the digit with aid of the counterweight. Each digit produced a unique force trace, and we extracted three metrics to allow cross-digit comparison: (i) initial slope, (ii) peak load, and (iii) average work over the first 2.5 mm following peak load (**Figure 3C**). We performed testing prior to surgery, post-injury and repair without added material, and following PTw application. Force traces from a single digit under all conditions showed tendon injury and repair altered the initial load onset, the excursion distance the digit began to flex, possibly reflecting a slight shortening of the tendon due to the repair (**Figure 3D**). Load slope was not significantly impacted by injury or material, but peak load and average work were significantly reduced following injury (*p*=0.018 and 0.024). PTw hydrogel did not significantly change peak load nor average work compared to injury without material (*p*=0.67 and 0.46) (**Figure 3E -G**). Following repeated cycles of flexion and extension on the MTS, incisions in each digit were reopened to visually evaluate the PTw material. It was visually observed that the PTw hydrogel remained at the application site, coating the tendon and sutures as well as between the FDP and FDS tendons, as highlighted by white arrows in **Figure 3A (v)**. We additionally subjected PNP hydrogel to the same testing in cadaver digits and found it performed similarly, with no impact on tendon peak load, slope, or average work compared to injury only, and did not reduce the stiffness or elasticity of the tendons (**Figure S4**).

**Figure 3.**
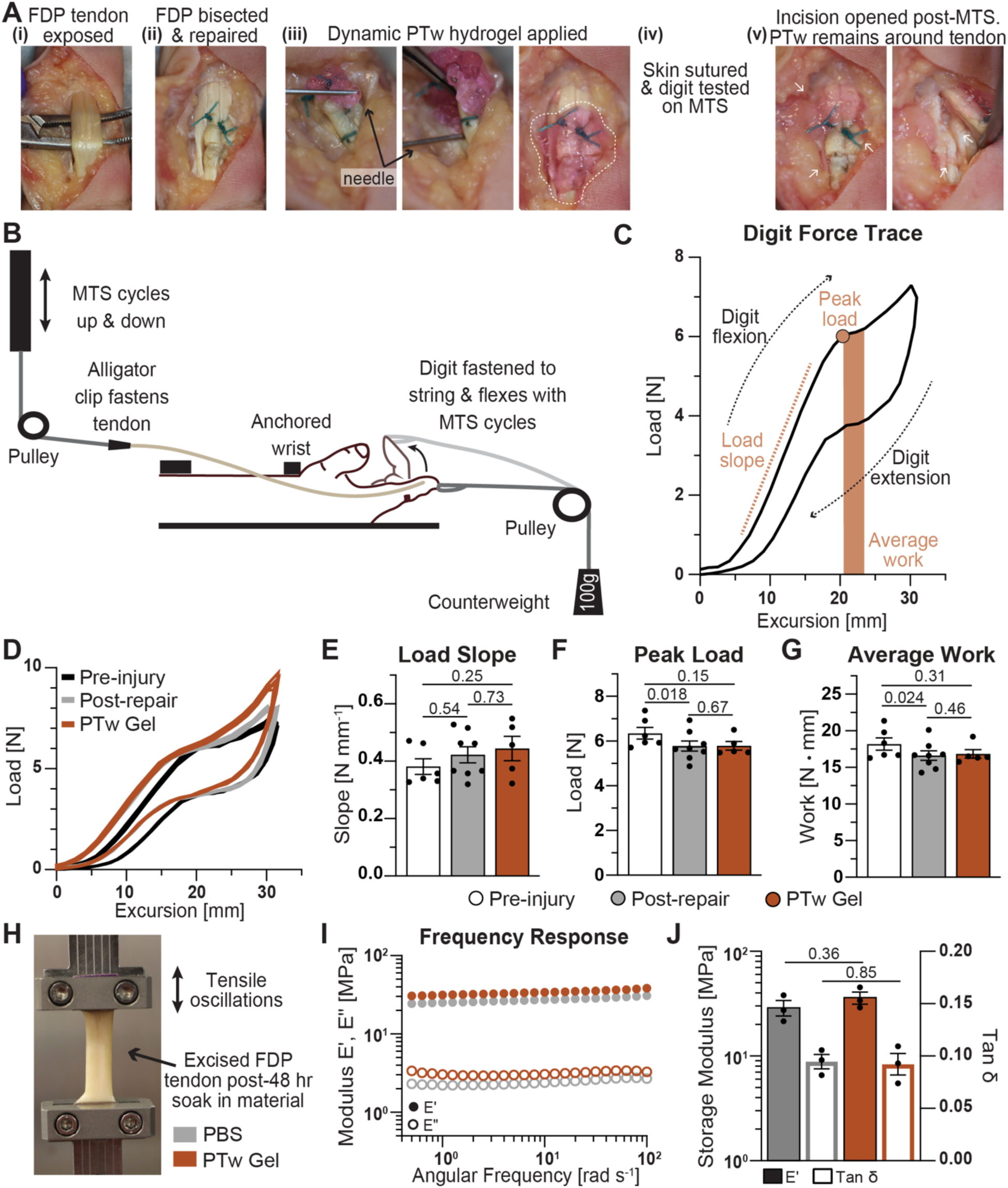
PTw application in human cadaver hands. A) Photos of (i-ii) tenotomy with repair followed by (iii) PTw hydrogel application, (iv) mechanical testing, and (v) evaluation of PTw retention around FDP. PTw hydrogel contains rhodamine-tagged HPMC-C_18_ for visualization and is denoted by dashed outline in (iii) and white arrows to regions with more difficult to see material in (v). B) Schematic of MTS setup with human cadaver arm. C) A single force trace for an uninjured ring digit showing curve direction as the digit is flexed and extended, as well as extracted parameters for load slope, peak load, and average work. D) Force traces of five replicate tests at 15 mm s^-1^ for a ring digit pre-injury (black or open bar), post-repair but before PTw application (grey), and after PTw application (orange). E) Extracted load slope, F) peak load, and G) average work across digits. Data presented as mean ± SEM, n = 5 - 8 across twelve unique digits from four arms and two patients. Statistical values shown are *p* values obtained from GLM fitting and Tukey HSD multiple comparison test in JMP (blocking by digit). H) Photo of excised FDP tendon mounted for DMA after 48 hrs soaking in PBS (grey) or PTw hydrogel (orange). I) Full frequency sweep and J) extracted values of E′ and tan δ at 40 rad·s^-1^ showing relative stiffness and elasticity of tendons is not impacted by hydrogel material compared with PBS. Data shown as mean ± SEM, n=3, statistical values are *p* values obtained from unpaired, two-tailed t-tests performed in GraphPad Prism.

In addition to showing PTw did not impair normal digit motion, we investigated the impact of prolonged exposure to PTw material on tendon integrity. Following excision, we soaked FDP tendon sections in PBS-soaked gauze or PTw hydrogel for 48 hours at 4 °C and assessed tendon mechanical properties by dynamic mechanical analysis (DMA) (**Figure 3H**). The frequency response of tendons remained the same for PBS or PTw soaked samples, with no significant difference in stiffness, as measured by storage modulus E′, nor elasticity represented by tan δ (**Figure 3I, J**). These studies demonstrate that PTw hydrogel is simple for a surgeon to apply with precision in the limited space of the digit, remains at the site of application following repeated cycles of digit flexion and extension, and does not impair meaningful metrics of digit motion like peak force or average work. Furthermore, PTw material does not impact tendon integrity following prolonged exposure times.

### 2.4 Evaluation of PTw in preclinical rodent Achilles tendon repair model

Having demonstrated key mechanical properties of PTw hydrogels which enable them to act as lubricious physical barriers when applied to human tendons without impeding gliding, we next evaluated the local retention and efficacy of PTw hydrogels in a preclinical rodent tendon injury model. We performed tenotomies on the left leg of Sprague-Dawley rats (N=24) wherein the Achilles tendon was completely divided and repaired with a standard modified Kessler suture. Following repair, eight control animals received no additional treatment prior to skin closure in accordance with the current clinical standard, and eight animals had PTw applied atop and around the tendon surgical site prior to skin closure. An additional eight animals were treated with PNP hydrogel in the same fashion as PTw hydrogel. To track retention of the material over time, PTw hydrogel was formulated with a NIR dye tagged HPMC-C_18_ and animals were imaged using in vivo imaging system (IVIS) for three weeks. Functional metrics of recovery were assessed with a catwalk and video gait analysis at week one prior to surgery and weeks one and eight following surgery. Animals were euthanized at week eight following surgery to evaluate hydrogel impacts on tendon healing (**Figure 4A**). Due to its facile injectability and tissue adhesive properties, the PTw hydrogel was easy to apply with precision in the small space of the rat ankle and remained adherent to the tissues during gentle spreading, ensuring all surfaces were covered (**Figure 4B**).

**Figure 4.**
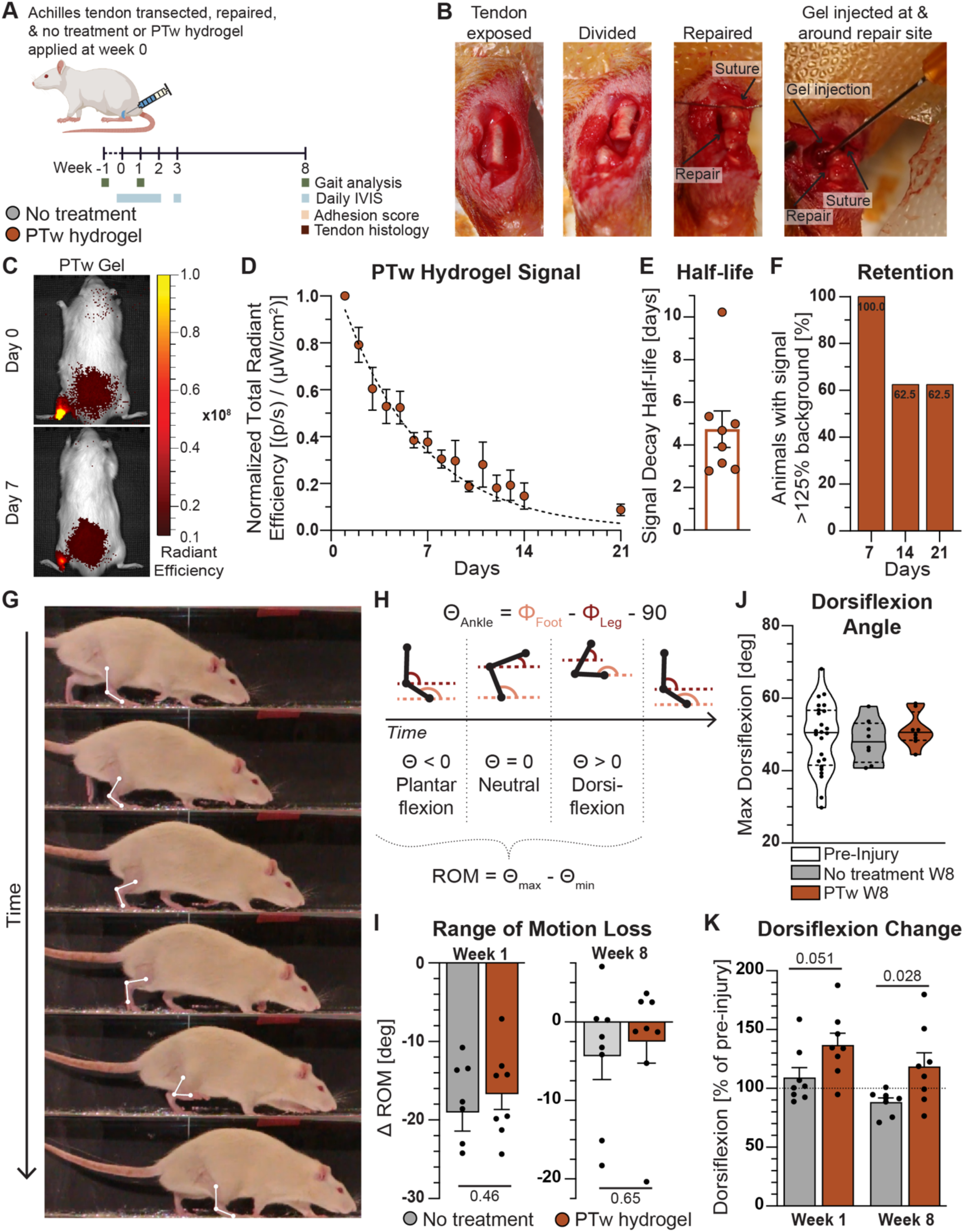
PTw retention and impact on rat Achilles tendon injury recovery. A) Timeline for rat Achilles tendon injury experiments involving tendon injury and repair followed by application of no treatment as a clinical standard control or PTw hydrogel and assessment of hydrogel retention via IVIS imaging and rat functional recovery with gait analysis experiments. B) Photos of surgical transection and repair of rat Achilles tendon followed by simple application of PTw hydrogel via injection and spreading before skin closure. C) Representative images of rats with PTw hydrogel on days zero and seven following surgery. D) Normalized fluorescent signal plotted over time along with E) extracted half-lives and F) fraction of animals with signal at least 125% of background animal signal. Total radiant efficiency of equal sized regions of interest was normalized to day 1 signal and half-lives extracted by fitting one-phase exponential decay in GraphPad Prism. Data in E shown as half-life for individual animals along with mean ± SEM. G) Photos showing a rat’s single stride and the vectors and angle from the knee to toes. H) Schematic explanation of the ankle angle, plantar or dorsiflexion, and range of motion (ROM). I) ROM difference in degrees for each rat at weeks one and eight compared to pre-injury. J) Dorsiflexion angle for all rats pre-injury and those in no treatment or PTw hydrogel groups at week eight post-injury. K) Fractional change in the dorsiflexion angle for rats at weeks one and eight normalized to pre-injury. Data presented as mean ± SEM, n=8. Statistical values are *p* values obtained from unpaired, two-tailed t-tests performed in GraphPad Prism.

We first characterized local retention of PTw hydrogel in the rat ankle and evaluated if the material remained present during crucial early stages of healing which are dominated by inflammation and high likelihood of adhesion initiation.^1,25,26^ Daily IVIS imaging for two weeks and once more at week three revealed a strong fluorescent signal at the application site even after one week (**Figure 4C**). We fit normalized fluorescence for each animal with an exponential decay and found the average half-life of PTw material to be 4.7 days, covering the key early inflammatory stage of healing (**Figure 4D, E**). Further, PTw-treated animals retained significant fluorescence signal throughout the entire three-week period, with more than half of the animals showing signals greater than 25% above the background from control animals with no hydrogel treatment at weeks two and three (5 in 8 animals) (**Figure 4F**). We additionally gauged retention of PNP hydrogel material and found a similar half-life of 5.5 days (**Figure S5**).

We next examined how PTw treatment impacted crucial recovery metrics by evaluating how each rat walked before injury and at weeks one and eight after injury using a catwalk setup and video gait analysis. We used three markers on each rat as reference points for image processing (the proximal point of the tibia or the lateral epicondyle, the calcaneus, and the fifth metatarsal head) and extracted the ankle angle throughout each gait. We defined the ankle angle according to A. S. P. Varejão and coworkers, where a value of 0° marked a neutral position of 90° between the leg and foot, negative values represented obtuse angles or plantar flexion, and positive values indicated acute angles or dorsiflexion (**Figure 4G, H**).^34^ One week after surgery we saw a substantial loss of range of motion (ROM) for all rats, with an average loss of -19.0° for control animals which received no treatment and a mitigated average loss of -16.7° for PTw treated animals. By week eight, the reduction in ROM was lower as the animals healed and recovered, but the no treatment animals maintained a greater average loss at -4.4° while PTw animals continued to show mitigated loss at an average of only -2.5° (**Figure 4I**). Given dorsiflexion represents a state of high strain for the Achilles tendon as it is stretched to bring the toes closer to the knee, we evaluated changes in the dorsiflexion angles for animals. The median dorsiflexion angle in no treatment animals at week eight was lower than the median of all animals pre-injury by -2.5°, indicating reduced dorsiflexion capacity, while the median angle for PTw animals was equivalent to pre-injury around 50.5°. Additionally, the spread of dorsiflexion angles represented by the interquartile range (IQR) was almost twice as large for no treatment animals at week eight than for PTw animals (7.5° compared to 4.1°), indicating PTw treatment contributed to more consistent recovery of dorsiflexion across animals (**Figure 4J, Table S1**). We also examined the ratio of dorsiflexion angles for each animal at weeks one and eight relative to pre-injury (**Figure 4K)**. No treatment and PTw treated animals both showed increases in dorsiflexion at week one of 9% and 37% on average, but only PTw treated animals had significantly greater dorsiflexion than pre-injury (*p*=0.0004, **Table S2**), likely due to the lubricious nature of the PTw hydrogel treatment. By week eight, dorsiflexion worsened in the control animals to lower than pre-injury values, with an average loss of -12% (*p*=0.086) compared to pre-injury and a three-fold loss of -21% from week one, likely due to adhesion formation and maturation limiting the Achilles tendon’s motion and stretch in dorsiflexion. On the other hand, PTw treated animals maintained increases in dorsiflexion at week eight, with an average increase of 19% compared to pre-injury (*p*=0.10), a significant difference compared to no treatment (*p*=0.028, **Figure 4K**). We also assessed these functional metrics for PNP hydrogel treated animals and found a similar trend wherein PNP material allowed for improved dorsiflexion immediately following surgery at week one (by 19%), but this benefit was reduced by week eight with PNP treated animals showing only a slight improvement in dorsiflexion angle over no treatment animals with a loss of -10% compared to -12% (**Figure S5D**). These studies showed PTw hydrogel treatment could be applied with precision in a preclinical tendon injury model and be retained at the application site for critical healing time frames, mitigating losses in key functional readouts like ROM and dorsiflexion.

### 2.5 Impact of PTw on healing of rodent Achilles tendon

Following functional evaluation of PTw hydrogel in rodent Achilles tendon injury, we further investigated PTw hydrogel biocompatibility in this peritendinous application. We carefully harvested the Achilles tendons of rats eight weeks after surgery and assessed the tendon repair strength and healing (**Figure 5A**). The original surgical suture used to repair the Achilles tendon was visible upon harvest and injured tendons were typically covered in adhesions which connected to the hypodermis as well as the fascia and muscle within the leg, regardless of treatment group. Consistent with other Achilles injury studies in rats, injured tendons tended to have a larger cross-sectional area than uninjured tendons due to generation of adhesion tissue around the repair (**Figure S6**).^35,36^ We mechanically characterized the elastic properties and ultimate strength of injured and uninjured tendons using tensile testing. Tendon samples were mounted with the calcaneus set at a 90° angle to the tendon-gastrocnemius insertion to better recapitulate physiological loading of the Achilles tendon at the tendon-bone interface (TBI) and pulled to failure (**Figure 5B**). Uninjured tendons consistently had higher strength and modulus than injured tendons, as would be expected after complete transection and repair (**Figure 5C-E**). PTw hydrogel treatment following injury did not have a significant impact on the strength nor modulus of healed tendons, demonstrating PTw hydrogel did not impair healing or reduce tendon mechanical integrity beyond reductions resulting from the surgery itself (**Figure 5D-E**). We additionally evaluated tendons treated with PNP hydrogel and similarly found no reduction in mechanical properties, indicating comparable healing to untreated controls (**Figure S7**).

**Figure 5.**
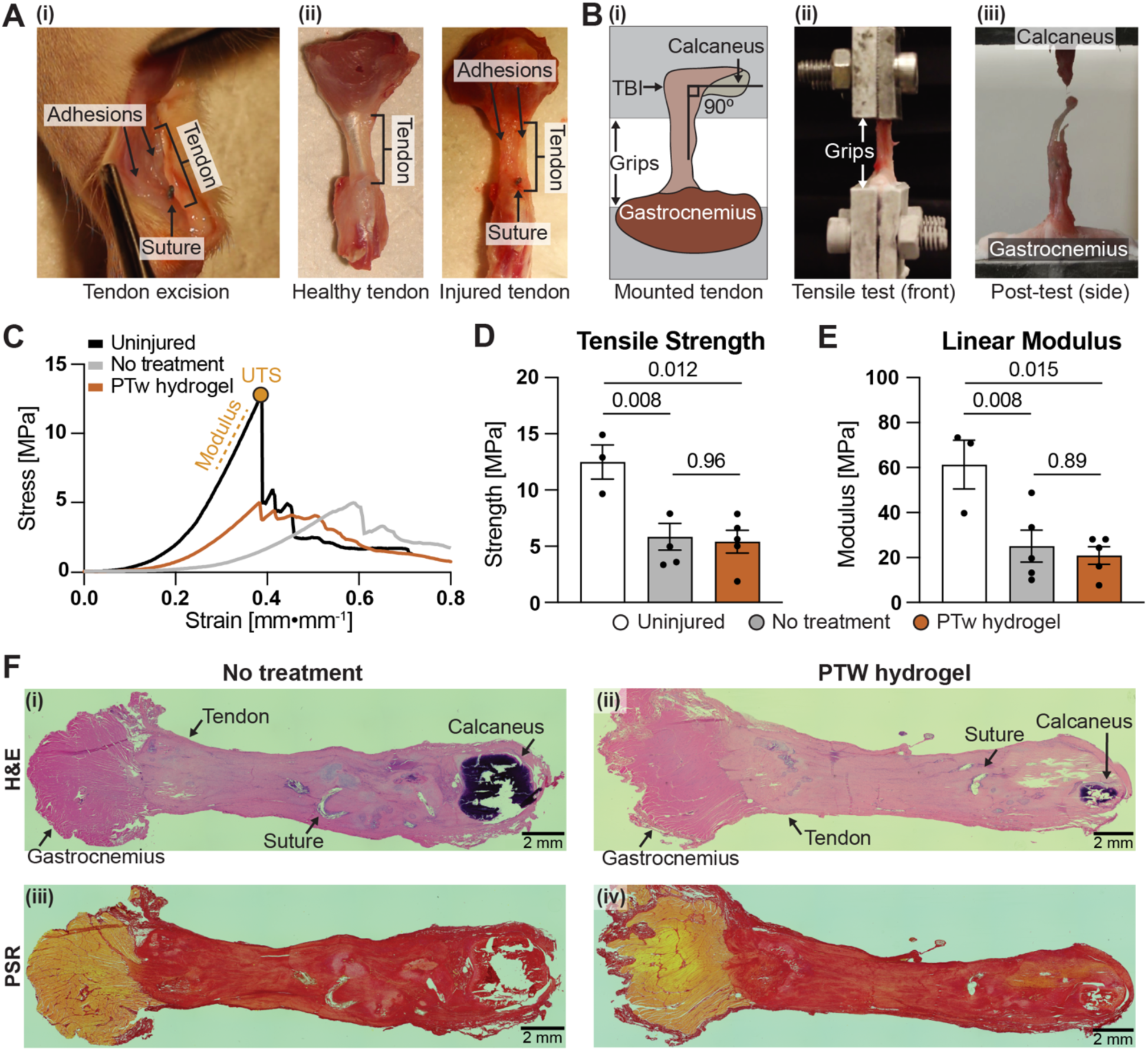
Harvest and mechanical testing of injured rat tendons. A) Excision of rat tendon (i) during adhesion scoring and representative images of (ii) healthy and injured tendons with adhesions labeled. B) Tensile testing of tendon with i) schematic of mounting, (ii) front view of tendon during testing, and (iii) side view after testing. C) Representative stress-strain curves for uninjured and injured tendons with relevant metrics represented schematically. D) Ultimate tensile strength and E) linear modulus of uninjured tendons and injured tendons. Data presented as mean ± SEM, n=3 for uninjured group, else n=5. Statistical values are *p* values obtained using GLM with Tukey’s test performed in JMP. F) Representative histology images (2.5x zoom) of injured tendons in each treatment group at week 8 with H&E staining (i, ii) and picrosirius red (iii, iv) showing no observable differences in healing.

We further evaluated the adhesions and healing of tendons with gross adhesion scoring and histological examination of hematoxylin and eosin (H&E) and picrosirius red (PSR) stained sections. For both hydrogels, gross adhesion scores were not severe (< 4) and not significantly different from untreated tendons (**Figure S8**). Blinded histological analysis by a pathologist indicated tendons in all treatment groups were well healed by week eight and there were no observable differences among groups (n=3 tendons per group) (**Figure 5F**). There was evidence of chondroid and osseus metaplasia in injured tendons; however, this was found in all samples and unrelated to PTw or PNP hydrogel material (**Figure S9**). Inflammation observed in H&E sections was limited to suture sites, with no indications of inflammation near tendon edges where there was contact with hydrogel material. In summary, these studies demonstrated PTw hydrogel as a safe, clinically effective, easy-to-apply treatment that reduced the impact of post-operative adhesions in a pre-clinical tendon surgery model and did not impar tendon healing relative to standard of care.

## 3. Discussion

In considering a biomaterial solution to peritendinous adhesions, several key desirable traits should be targeted: (i) simple application during surgery, (ii) retention at application site throughout the critical early inflammatory and proliferative time frames, (iii) improvement of clinically relevant functional outcomes such as ROM, and (iv) safety, showing no negative impact on tendon healing or repair strength. In this work we present a dynamic hydrogel, PTw, which meets these criteria and shows promise as an anti-adhesion intervention following tendon surgery. PTw mechanical properties were invariant to temperature, ensuring the material behaved the same during prolonged refrigerated storage, room temperature application, and *in vivo* usage. Shear and extensional rheology elucidated the tissue-adhesive properties of PTw hydrogels which ensured the material adhered to relevant tissues following application but allowed tissues to easily slide past one another with bulk material yielding. These properties are crucial for application in the limited and highly mobile space surrounding the digital flexor tendon, where the tendon must glide freely through pulleys and the applied material must remain at the application site while withstanding the strains of that motion. Indeed, PTw hydrogel formed a lubricious barrier which enabled early mobilization following surgery, an important rehabilitation regiment aimed at reducing adhesions.^14,15^ We confirmed in human cadaver digits following clinically relevant injury and repair to zone II FDP tendons that PTw application did not impair tendon glide nor reduce tendon strength and integrity after prolonged contact. We next determined the efficacy and safety of PTw hydrogels in a preclinical rat Achilles tendon injury model and found improved range of motion and significantly improved dorsiflexion compared to control animals following eight weeks of healing. PTw material was simple to apply and remained at the repair site for over three weeks, covering critical stages in tendon healing that may lead to adhesion formation. PTw hydrogel was also safe and did not impair tendon healing nor strength. Overall, this work reports a novel solution to the problem of peritendinous adhesions and elucidates the viscoelastic and flow properties that make PTw hydrogel suited for this application.

Hydrogels have been increasingly explored as materials to limit adhesions following tendon surgery owing to their tunability and biocompatibility.^37-43^ Investigated materials range widely in composition and utilize synthetic components like Prussian blue nanoparticles and blood-derived platelet lysate, chitosan and hyaluronic acid-grafted poly(N-isopropylacrylamide) (PNIPAM), or polyglycerol-grafted graphene and molybdenum disulfide.^37,41,42^ While the diversity of hydrogel materials being explored for this application is encouraging, many are complex and bespoke, making a path to good manufacturing practice (GMP) and clinical approval difficult. In addition to hydrogels, electrospun fibers are another material modality heavily explored in the literature.^9,44^ Unfortunately, there remain to date no materials on the market with FDA approval to prevent peritendinous adhesions, due in part to the cost and complexity of manufacturing for many materials, including electrospun fibers.^9,44^ PTw hydrogels, in contrast, are composed of two simple and commercially available components, a cellulose derivative and tween 20, both of which are produced at large scales as cosmetic additives and drug excipients.^45,46^

Another major drawback of many preclinical animal studies of peritendinous adhesions is a lack of evidence for functional recovery. Many studies primarily measure the gross incidence of adhesions upon visual observation, but this metric is subjective and imprecise compared to a quantitative measure of function recovery.^47^ Additionally, while there is currently no regulatory guidance from the FDA on how to best design or execute clinical trials for anti-adhesion agents, the assessment of adhesion score by non-invasive means should complement patient functional outcome metrics, like ROM.^48^ Interestingly, in this work we found clear improvement in functional outcomes but no significant change in adhesion incidence scores with PTw or PNP hydrogel treatment (**Figure 4**, **Figure S8**). Here it should be noted that the Achilles tendon of a rat does not possess the same synovial sheath anatomy as a human flexor tendon or the flexor tendons of larger animals such as turkeys, rabbits, or dogs.^16,41,47,49-52^ Though, the rat Achilles injury model is well suited for assessment of functional recovery over time as well as investigating the safety profile of hydrogels.^53,54^ Future studies should be conducted in larger animal models focused on assessing gross adhesion severity along with functional recovery, as well as modulating the formulation of PTw to optimize its mechanical properties for this application.

It is likely that a successful platform for preventing peritendinous adhesions following surgery will combine a physical barrier with delivery of molecules to modulate the inflammatory and cellular responses.^9,44^ We have previously shown that dynamic hydrogels including and similar to PTw and PNP are amenable to long term delivery of various cargos, including small molecules and proteins.^46,55-57^ Future work could expand the PTw platform for sustained local delivery of small molecule modulators of inflammation, like corticosteroids or ibuprofen to reduce initial inflammation, or protein therapeutics like vascular endothelial growth factor (VEGF) or other growth factors to promote healing, while acting as a lubricious barrier.^58-63^

## 4. Materials and Methods

*Hydrogel formulations:* PTw hydrogels were prepared as previously described.^24^ HPMC-C_18_ (Sangelose 90L, Daido chemical corp.) was dissolved in phosphate buffered saline (PBS, pH 7.4) at 4 wt% and loaded into a luer-lock syringe. Tween 20 (Sigma-Aldrich) was diluted in PBS to a concentration of 10% v/v, then diluted further for each hydrogel prepared such that the final concentration in the hydrogel was 0.5%. This Tween 20 - PBS solution was loaded into a second syringe and the two syringes were connected with a female-female luer lock elbow with care to avoid air at the interface. The solutions were mixed until a homogenous PTw hydrogel was formed. All PTw hydrogels were formulated at 1.8 wt% HPMC-C_18_ and 0.5% Tween 20. Hydrogels were stored at 4 °C until use. PNP hydrogels were prepared as previously reported and PNP-1-10 (1 wt% HPMC-C_12_ and 10 wt% PEG-PLA NPs) was the only formulation used in this work.^64^

*Fluorescent hydrogel formulations:* HPMC-C_18_ and C_12_ were fluorescently tagged with NIR and rhodamine for visualization in cadaver experiments and IVIS imaging in rats. First, 1 g HPMC-C_x_ was dissolved in NMP (40 mL) at room temperature with stirring. Once the polymer had completely dissolved, the reaction was brought to 50 °C and a solution of either NIR-797 isothiocyanate or rhodamine B isothiocyanate (Santa Cruz Biotechnology, 1 mg / mL or 1.14 mmol and 1.86 mmol respectively) in NMP (10 mL) was added dropwise, followed by DIPEA (catalyst, 125 µL). The reaction was maintained at 50 °C for 30 minutes, then heat was shut off and mixture was left stirring overnight at room temp. The solution was then precipitated from acetone and polymer-NIR 797 or -rhodamine was purified by dialysis against MilliQ water for 4 - 7 days (MWCO 3.5 kDa) and lyophilized, yielding polymer as a green (NIR) or pink (rhodamine) powder. Each polymer was dissolved in sterile PBS, pH 7.4, prior to use in hydrogels. For cadaver application, PTw hydrogels were formulated with 1.8 wt% HPMC-C_18_-rhodamine and PNP with 1 wt% HPMC-C_12_-rhodamine. For rat studies, PTw hydrogels were formulated with 1.8 wt% HPMC-C_18_-NIR 797 and PNP with 1 wt% HPMC-C_12_-NIR 797.

*Hydrogel rheological characterization:* Rheological characterization was performed on hydrogels using a TA Instruments DHR-2 stress-controlled rheometer. All experiments were performed using a 20 mm diameter serrated plate geometry at 25 °C with a 500 µm gap. Flow sweeps were performed from high to low shear rates. Step shear experiments were performed by alternating between a low shear rate (0.1 s^−1^; 60 s) and a high shear rate (10 s^−1^; 60 s) for three cycles. Frequency sweep measurements were performed at a constant 1% strain in the linear viscoelastic regime. Amplitude sweeps were performed 10 rad s^-1^. Values for storage modulus and tan δ were reported from frequency sweeps at the 40 rad s^-1^ frequency. Yield strain was defined as the first strain value after G′ < 0.85 * maximal pre-yield G′ value. For temperature controlled experiments, temperature was set to 4, 25, or 37 °C.

*Tissue adhesion characterization:* Stress sweeps were performed from low to high with steady state sensing and yield stress was defined as the first stress value after viscosity < 0.85 * maximal pre-yield viscosity value. For control runs, two 20 mm serrated parallel plates were used. For tissue runs, cadaver skin, murine skin, or cadaver tendon tissue was positioned in place of the bottom rheometer plate. Before loading hydrogels onto cadaver skin, sutures were sewn into the edges of the sample to act as anchors, which were then taped down to flatten the tissue under the geometry. Kimwipes were used to remove excess fluid from the surface of tissue samples and fatty protrusions were removed with a scalpel to further flatten the tissue surface. For tendon tissue, multiple tendons were arranged with minimal gapping and as flat a surface as possible. All experiments were performed at 25 °C with a 500 µm gap.

*Extensional rheology:* Filament stretching extensional rheometry was performed on a TA Instruments ARES-G2 in axial mode using an 8 mm plate geometry starting with a 4 mm gap to test an initial radius-to-height aspect ratio of 1:1. All experiments were conducted at 25 °C with a Hencky (exponential) strain rate of 0.1 s^-1^. Experiments were recorded with a DSLR camera and the peak extension of the hydrogel was measured using Tracker 6.1.3 software (Open Source Physics). The reported strain is the ratio of extension at break to the initial length of 4 mm.

*MTS studies on cadaver arms:* A custom rig was engineered to secure cadaver arms for testing glide friction of tendons following repair and hydrogel application. To simulate injury, flexor digitorum profundus (FDP) tendons in the index, middle, and ring digits were isolated and severed between the A2 and A4 pulleys (n=12 digits, 4 arms, 2 individuals). Tendons were then repaired with a modified Kesler knot using 4-0 polypropylene sutures (Ethicon) and the skin sutured closed. In treatment groups, 200 – 300 μL hydrogel was injected on top of and around the repaired tendon and manipulated to coat the tendon before the skin was closed. An MTS Bionix 200 test system was used to apply displacement and measure force. The cadaver arm was positioned atop an optical table (Thorlabs) and secured with two S-brackets (forming an arch) and bolts from the S-brackets to the pisiform and scaphoid wrist bones. An incision was made in the forearm to access the FDP tendons, which were severed at the proximal end, wrapped in sandpaper, and secured by an alligator clip to a nylon string fed through a pulley and into the MTS clamp. Nylon string was also tied to a suture loop on each fingertip, passed over a pulley, and attached to a 100 g counterweight to fully extend the digit being tested and allow full range of motion during glide experiments. Force was tared and slack removed from the string and tendon before each experiment. Experiments were conducted to an excursion of 35 - 40 mm at speeds of 5 and 15 mm s^-1^. Bolts, brackets, nuts, alligator clips, and string were purchased from McMaster-Carr.

*MTS data processing:* Force vs. displacement traces were analyzed using custom code in MATLAB (MathWorks, Natick, MA). The load slope was calculated as the slope of the line from 0.5 N of loading to 4.5 N of loading for each curve. The peak load was defined as either the first local maximum of the loading curve or, in cases where the curve did not have a local maximum before unloading, the point on the curve at which the slope decreased to 40% of the load slope. The excursion corresponding to the peak load was determined to be the start of the plateau region. Work of flexion was defined as the area under the curve to an excursion of 2.5 mm past the start of the plateau region. All extracted metrics are the average of 9-10 loading and unloading cycles.

*DMA on cadaver tendons:* After MTS studies were performed, uninjured 4 cm segments of FDP tendons were secured from all digits and either wrapped in PBS-soaked gauze or immersed in PNP or PTw hydrogels for 48 hours at 4 °C. Tendons were mounted on a TA Instruments ARES-G2 strain-controlled rheometer-DMA with 150 grit sandpaper (3M Company) to prevent slip within the grips. Samples were pre-loaded to 0.1 N before testing and frequency sweeps were performed at an oscillation strain of 0.1%. Values for storage modulus and tan δ were reported from frequency sweeps at 40 rad s^-1^.

*Rat Achilles tendon injury and repair model:* All animal studies were performed in accordance with the National Institutes of Health (NIH) guidelines, with the approval of the Stanford Administrative Panel on Laboratory Animal Care. Adult male Sprague-Dawley rats (450-600 g, 15-17 weeks, n=24) were purchased from Charles River and housed in the animal facility at Stanford University. Animals were randomly assigned to treatment groups (n=8 each) and housed such that each cage had two animals with different treatments to mitigate cage effects. For surgery, animals were anesthetized under 3.5% (v/v) isoflurane, their left hind leg shaved and aseptically prepared, and a longitudinal incision made to expose the Achilles tendon. The accessory Achilles was first isolated and removed (∼8 mm), then the Achilles tendon was isolated, transected approximately 5 mm above its insertion site into the calcaneus, and repaired with 5-0 ethibond sutures (Ethicon) using a modified Kessler suture. Following repair, approximately 100 μL hydrogel was applied to the repair site and manipulated to fully coat the repaired tendon in treatment animals (n=8 with PTw and n=8 with PNP). Skin was then sutured closed with 5-0 nylon sutures (Ethicon) and animals were returned to paired housing without leg immobilization. Animal gait analysis was performed at week one prior to surgery and again at weeks 1 and 8 following surgery. IVIS imaging was performed immediately following surgery and then daily for two weeks with a final time point at three weeks. Animals were euthanized eight weeks following surgery, to allow for sufficient healing and adhesion maturation, and their injured leg dissected and tissues harvested for histological processing (n=3 each treatment) and mechanical testing (n=5 each treatment). Contralateral uninjured tendons were also harvested for controls from randomized animals.

*IVIS imaging:* Rats were imaged using an IVIS (model, Lago) daily for two weeks and again at week three. Rats were anesthetized with 2.5% (v/v) isoflurane and imaged with auto exposure settings at an excitation wavelength of 780 nm and emission wavelength of 845 nm. Adaptive fluorescent background subtraction was applied and total radiant efficiency ([p/s] / [µW/cm²]) was quantified using an equal-sized region of interest around the left leg for each rat (the injury site). This was also done for rats that did not receive hydrogel material to provide an average background signal which was subtracted from the signal for rats that did receive hydrogels. Because some animals had fluorescence increase between day 0 and 1, the background-subtracted total radiant efficiency at each time point was normalized to that animals’ signal on day 1. Normalized values were then fit to a one-phase exponential decay and half-lives calculated for each animal in GraphPad Prism.

*Functional recovery following rat tendon injury:* A transparent walkway (91 cm x 14 cm x 10 cm) was constructed from plexiglass to record the gait of rats for this study. Before recording, rats were anesthetized with 2.5% (v/v) isoflurane, their left hind limb shaved, and three points marked with red sharpie for video analysis: the proximal tibia, calcaneus, and fifth metatarsal head. Rats were placed at one end of the walkway and their home cage placed at the other end to encourage walking. At each time point, each animal was recorded walking with a DSLR camera at 60 frames per second and a shutter speed of 2000 s^-1^. A walk was considered successful if at least three consecutive steps were taken. At least two steps were analyzed for each rat. All data were processed first by marking the proximal tibia, calcaneus, and fifth metatarsal positions with Tracker 6.1.3 software. The positions of these points were then processed using custom MATLAB code to identify the foot and leg angles and calculate the ankle angle. The clockwise leg angle from horizontal was subtracted by the clockwise foot angle from horizontal and 90° was further subtracted from this to set a neutral angle of 0° when the foot and leg segments made a 90° angle. A rat’s leg was defined as dorsiflexed when the ankle angle was greater than 0° and plantarflexed when the ankle angle was less than 0°. Active range of motion was calculated by the difference between maximum dorsiflexion and plantarflexion angles.

*Mechanical testing of rat tendon:* Following tendon injury, repair, and eight weeks of healing, the Achilles tendons of Sprague-Dawley rats were excised by cutting at the gastrocnemius and fine dissection of the calcaneus. Samples were refrigerated for 2-3 days before testing. The muscle end of the tendon was secured in the clamps after being coated with OCT Compound (Agar Scientific). The metal clamps were then dipped in liquid nitrogen, taking care not to submerge the tendon sample. After freezing, the frozen clamps were further tightened around the muscle. The calcaneus bone was secured within the clamps at a 90° angle to the direction of force, recapitulating the neutral angle of the leg. Tendon length was measured before mounting into the grips and samples were (1) preloaded to 0.1 N; (2) measured using calipers; (3) subjected to 10 cycles of preconditioning to 2% strain; and (4) pulled to failure. The strain rate was set to 4% s^-1^ for all steps. All samples failed at the tendon-bone interface. The linear modulus was calculated by extracting the slope of the resultant stress-strain curves in the linear regime and the ultimate tensile strength was defined as the peak of the stress-strain curve.

*Tendon histology:* Following tendon injury, repair, and eight weeks of healing, the Achilles tendons of nine Sprague-Dawley rats (three per treatment group) were excised by cutting at the gastrocnemius and fine dissection of the calcaneus. Samples were fixed in cassettes for 24 hrs in 10% neutral buffered formalin. Fixed samples were submitted to Histo-Tec Laboratory Inc. (Hayward, CA, USA) for decalcification, paraffin embedding, slicing (foot-based coronal plane), and H&E and PSR staining. Returned slides were imaged on a Leica LC221 THUNDER Imager using 2.5x and 10x objectives and individual fields of view were deconvoluted and stitched together to construct images of large tissue segments using built-in functionality of the LASX software.

*Statistics:* For in vivo experiments, animal treatments were randomized across cages and data is presented as mean ± SEM. Comparisons between multiple groups were conducted with the general linear model (GLM) and Tukey HSD test in JM P, accounting for digit blocking in the case of experiments on human cadaver hands. Comparisons between two groups were conducted with unpaired two-tailed t-tests run in GraphPad Prism. Select *p* values are shown in the text and figures and all *p* values are in the supporting information.

## Acknowledgements

The authors would like to thank every member of the Appel Lab, former and current, for their on-going support, technical expertise, and scientific discussion. In particular, the authors thank Noah Eckman and Samya Sen for their contributions to theorizing calculations, and Noah Eckman for assistance socializing animals. The authors would also like to thank Prof. Dauskardt and lab members for training on and use of their MTS instrument. Histological analysis was performed by Dr. José Vilches-Moure, DVM, PhD, with Stanford’s Veterinary Service Center Comparative Pathology services.

## Author contributions

Conceptualization: ELM, CMW, PMF, EAA

Methodology: ELM, CMW, YES, PMF

Software: CMW

Investigation: ELM, CMW, VMD, SJB, SCW, CKJ, PMF

Visualization: ELM, CMW Supervision: PMF, EAA

Writing—original draft: ELM, CMW

Writing—review & editing: SJB, SCW, PMF, EAA

## Data and materials availability

All data needed to evaluate the conclusions of this paper are present in the main text and/or the supplementary materials.

## References

1 Legrand, A., Kaufman, Y., Long, C. & Fox, P. M. Molecular Biology of Flexor Tendon Healing in Relation to Reduction of Tendon Adhesions. The Journal of Hand Surgery 42, 722–726, doi:10.1016/j.jhsa.2017.06.013 (2017).

2 Fenwick, S. A., Hazleman, B. L. & Riley, G. P. The vasculature and its role in the damaged and healing tendon. Arthritis Research 4, 252–260 (2002).

3 Daley, B. J., Cecil, W., Clarke, C. P., Cofer, J. B. & Guillamondegui, O. D. How Slow Is Too Slow? Correlation of Operative Time to Complications: An Analysis from the Tennessee Surgical Quality Collaborative. Journal of the American College of Surgeons 220, 550, doi:10.1016/j.jamcollsurg.2014.12.040 (2015).

4 Sikirica, V. et al. The inpatient burden of abdominal and gynecological adhesiolysis in the US. BMC surgery 11, 13, doi:10.1186/1471-2482-11-13 (2011).

5 Docheva, D., Müller, S. A., Majewski, M. & Evans, C. H. Biologics for tendon repair. Advanced Drug Delivery Reviews 84, 222–239, doi:10.1016/j.addr.2014.11.015 (2015).

6 Strickland, J. W. Flexor Tendon Injuries: I. Foundations of Treatment. JAAOS - Journal of the American Academy of Orthopaedic Surgeons 3, 44 (1995).

7 Titan, A. L., Foster, D. S., Chang, J. & Longaker, M. T. Flexor Tendon: Development, Healing, Adhesion Formation, and Contributing Growth Factors. Plastic and Reconstructive Surgery 144, 639e, doi:10.1097/PRS.0000000000006048 (2019).

8 Chinchalkar, S. J., Larocerie-Salgado, J. & Suh, N. Pathomechanics and Management of Secondary Complications Associated with Tendon Adhesions Following Flexor Tendon Repair in Zone II. Journal of Hand and Microsurgery 8, 70–79, doi:10.1055/s-0036-1586173 (2016).

9 Zhang, Q., Yang, Y., Yildirimer, L., Xu, T. & Zhao, X. Advanced technology-driven therapeutic interventions for prevention of tendon adhesion: Design, intrinsic and extrinsic factor considerations. Acta Biomaterialia 124, 15–32, doi:10.1016/j.actbio.2021.01.027 (2021).

10 Hohendorff, B. et al. Tenolysis of extensor and flexor tendons of the hand. Der Orthopäde 49, 771–783, doi:10.1007/s00132-020-03965-x (2020).

11 Feldscher, S. B. & Schneider, L. H. FLEXOR TENOLYSIS. Hand Surgery, doi:10.1142/S0218810402000819 (2011).

12 Seppi, S., Vecchi, S., Raccagni, I., Novelli, C. & Pajardi, G. E. Pre- and post-treatment in flexor tendon tenolysis: An observational study. Journal of Hand Therapy, doi:10.1016/j.jht.2023.10.004 (2024).

13 Vinitpairot, C., Yik, J. H. N., Haudenschild, D. R., Szabo, R. M. & Bayne, C. O. Current trends in the prevention of adhesions after zone 2 flexor tendon repair. Journal of Orthopaedic Research **n/a**, doi:10.1002/jor.25874 (2024).

14 Starr, H. M., Snoddy, M., Hammond, K. E. & Seiler, J. G. Flexor Tendon Repair Rehabilitation Protocols: A Systematic Review. The Journal of Hand Surgery 38, 1712–1717.e1714, doi:10.1016/j.jhsa.2013.06.025 (2013).

15 Lister, G. D., Kleinert, H. E., Kutz, J. E. & Atasoy, E. Primary flexor tendon repair followed by immediate controlled mobilization. The Journal of Hand Surgery 2, 441–451, doi:10.1016/S0363-5023(77)80025-7 (1977).

16 Wan, R. et al. 109. Comparing the Postoperative Adhesion Prevention Effectiveness of Collagen-glycosaminoglycan (GAG), Hyaluronic Acid (HA), and Pentamidine Using a Turkey In Vivo Model. Plastic and Reconstructive Surgery – Global Open 11, 68, doi:10.1097/01.GOX.0000938024.46146.20 (2023).

17 Turner, J. B., Corazzini, R. L., Butler, T. J., Garlick, D. S. & Rinker, B. D. Evaluating Adhesion Reduction Efficacy of Type I/III Collagen Membrane and Collagen-GAG Resorbable Matrix in Primary Flexor Tendon Repair in a Chicken Model. HAND 10, 482–488, doi:10.1007/s11552-014-9715-x (2014).

18 McDermott, E. R., Bowers, Z. & Nuelle, J. A. The Application of Hyaluronic Acid/Alginate Sheet to Flexor Pollicis Longus Tendon Repair to Prevent Adhesion Formation: A Second Look. Cureus 14, e33147, doi:10.7759/cureus.33147 (2022).

19 McCahon, J. A. S., Bridges, T. N. & Parekh, S. G. Application of Hyaluronic Acid/Alginate Sheet to Achilles Tendon Injuries to Prevent Peritendinous Adhesions. Techniques in Foot & Ankle Surgery (2024).

20 Voleti, P. B., Buckley, M. R. & Soslowsky, L. J. Tendon Healing: Repair and Regeneration. Annual Review of Biomedical Engineering 14, 47–71, doi:10.1146/annurev-bioeng-071811-150122 (2012).

21 Appel, E. A. et al. Self-assembled hydrogels utilizing polymer–nanoparticle interactions. Nature Communications 6, 6295, doi:10.1038/ncomms7295 (2015).

22 Stapleton, L. M. et al. Use of a supramolecular polymeric hydrogel as an effective post-operative pericardial adhesion barrier. Nature Biomedical Engineering 3, 611–620, doi:10.1038/s41551-019-0442-z (2019).

23 Stapleton, L. M. et al. Dynamic Hydrogels for Prevention of Post-Operative Peritoneal Adhesions. Advanced Therapeutics 4, 2000242, doi:10.1002/adtp.202000242 (2021).

24 Song, Y. E., Eckman, N., Sen, S., Saouaf, O. M. & Appel, E. A. Highly extensible physically crosslinked hydrogels for high-speed 3D bioprinting. bioRxiv, 2024.2008.2005.606733, doi:10.1101/2024.08.05.606733 (2024).

25 Griffin, M., Hindocha, S., Jordan, D., Saleh, M. & Khan, W. Management of Extensor Tendon Injuries. Open Orthopaedics Journal, 36–42, doi:10.2174/1874325001206010036 (2012).

26 Pearce, O., Brown, M. T., Fraser, K. & Lancerotto, L. Flexor tendon injuries: Repair & Rehabilitation. Injury 52, 2053–2067, doi:10.1016/j.injury.2021.07.036 (2021).

27 Grimaldo Ruiz, O., et al. Finite element analysis of the flexor digitorum profundus tendon during a passive rehabilitation protocol. Revista Facultad de Ingeniería Universidad de Antioquia, 124–132, doi:10.17533/udea.redin.20210528 (2021).

28 Ugbolue, U. C., Hsu, W.-H., Goitz, R. J. & Li, Z.-M. Tendon and nerve displacement at the wrist during finger movements. Clinical Biomechanics 20, 50–56, doi:10.1016/j.clinbiomech.2004.08.006 (2005).

29 Sapienza, A., Yoon, H. K., Karia, R. & Lee, S. K. Flexor tendon excursion and load during passive and active simulated motion: a cadaver study. Journal of Hand Surgery (European Volume) 38, 964–971, doi:10.1177/1753193412469128 (2013).

30 Luotonen, O. I. V. et al. Benchmarking supramolecular adhesive behavior of nanocelluloses, cellulose derivatives and proteins. Carbohydrate Polymers 292, 119681, doi:10.1016/j.carbpol.2022.119681 (2022).

31 Tudoroiu, E.-E. et al. An Overview of Cellulose Derivatives-Based Dressings for Wound-Healing Management. Pharmaceuticals 14, 1215, doi:10.3390/ph14121215 (2021).

32 Stevens, K. A., Caruso, J. C., Fallahi, A.-K. M. & Patiño, J. M. in StatPearls (StatPearls Publishing, 2023).

33 McCarthy, D. M., Boardman, N. D., Tramaglini, D. M., Sotereanos, D. G. & Herndon, J. H. Clinical management of partially lacerated digital flexor tendons: A surgery of hand surgeons. The Journal of Hand Surgery 20, 273–275, doi:10.1016/S0363-5023(05)80023-1 (1995).

34 Varejão, A. S. P. et al. Motion of the foot and ankle during the stance phase in rats. Muscle & Nerve 26, 630–635, doi:10.1002/mus.10242 (2002).

35 Chamberlain, C. S. et al. Temporal Healing in Rat Achilles Tendon: Ultrasound Correlations. Annals of biomedical engineering 41, 477–487, doi:10.1007/s10439-012-0689-y (2013).

36 Huegel, J. et al. Quantitative Comparison of Three Rat Models of Achilles Tendon Injury: A Multidisciplinary Approach. Journal of biomechanics 88, 194–200, doi:10.1016/j.jbiomech.2019.03.029 (2019).

37 Wang, K., Chen, D., Wang, Z., Yang, J. & Liu, W. An Injectable and Antifouling Supramolecular Polymer Hydrogel with Microenvironment-Regulatory Function to Prevent Peritendinous Adhesion and Promote Tendon Repair. Macromolecular Bioscience **n/a**, 2300142, doi:10.1002/mabi.202300142 (2023).

38 Bassetto, F. et al. Efficacy and safety of Dynavisc® gel in prevention of scar adhesions recurrence after flexor tendons tenolysis in zone 2. Multicenter retrospective cohort study. Annali Italiani Di Chirurgia 94, 529–536 (2023).

39 von Kieseritzky, J., Rosengren, J. & Arner, M. Dynavisc as an Adhesion Barrier in Finger Phalangeal Plate Fixation—a Prospective Case Series of 8 Patients. Journal of Hand Surgery Global Online 2, 109–112, doi:10.1016/j.jhsg.2019.11.003 (2020).

40 Ishiyama, N. et al. The prevention of peritendinous adhesions by a phospholipid polymer hydrogel formed in situ by spontaneous intermolecular interactions. Biomaterials 31, 4009–4016, doi:10.1016/j.biomaterials.2010.01.100 (2010).

41 Chou, P.-Y. et al. Thermo-responsive in-situ forming hydrogels as barriers to prevent post-operative peritendinous adhesion. Acta Biomaterialia 63, 85–95, doi:10.1016/j.actbio.2017.09.010 (2017).

42 Barzegar, P. E. F. et al. Graphene-MoS2 polyfunctional hybrid hydrogels for the healing of transected Achilles tendon. Biomaterials Advances 137, 212820, doi:10.1016/j.bioadv.2022.212820 (2022).

43 Marchesini, A. et al. Effectiveness of Hyaluronan Autocross-Linked-Based Gel in the Prevention of Peritendinous Adherence Following Tenolysis. Applied Sciences 11, 7613, doi:10.3390/app11167613 (2021).

44 Zhou, H. & Lu, H. Advances in the Development of Anti-Adhesive Biomaterials for Tendon Repair Treatment. Tissue Engineering and Regenerative Medicine 18, 1–14, doi:10.1007/s13770-020-00300-5 (2021).

45 Aldawsari, H. et al. Combined use of cyclodextrins and hydroxypropylmethylcellulose stearoxy ether (Sangelose®) for the preparation of orally disintegrating tablets of type-2 antidiabetes agent glimepiride. Journal of Inclusion Phenomena and Macrocyclic Chemistry 80, 61–67, doi:10.1007/s10847-014-0386-6 (2014).

46 Liang, Y.-K., Cheng, W.-T., Chen, L.-C., Sheu, M.-T. & Lin, H.-L. Development of a Swellable and Floating Gastroretentive Drug Delivery System (sfGRDDS) of Ciprofloxacin Hydrochloride. Pharmaceutics 15, 1428, doi:10.3390/pharmaceutics15051428 (2023).

47 Chang, J., Thunder, R., Most, D., Longaker, M. T. & Lineaweaver, W. C. Studies in Flexor Tendon Wound Healing: Neutralizing Antibody to TGF-β1 Increases Postoperative Range of Motion. Plastic and Reconstructive Surgery 105, 148 (2000).

48 De Wilde, R. L. et al. The Future of Adhesion Prophylaxis Trials in Abdominal Surgery: An Expert Global Consensus. Journal of Clinical Medicine 11, 1476, doi:10.3390/jcm11061476 (2022).

49 Potenza, A. D. Tendon Healing Within the Flexor Digital Sheath in the Dog: An Experimental Study. JBJS 44, 49 (1962).

50 Boz, M. et al. Does methylene blue reduce adhesion during the healing process after tendon repair? Joint Diseases and Related Surgery 31, 246–254, doi:10.5606/ehc.2020.74405 (2020).

51 Zhao, C. et al. Effects of a Lubricin-Containing Compound on the Results of Flexor Tendon Repair in a Canine Model in Vivo. JBJS 92, 1453, doi:10.2106/JBJS.I.00765 (2010).

52 Karakurum, G., Buyukbebeci, O., Kalender, M. & Gulec, A. Seprafilm® interposition for preventing adhesion formation after tenolysis: An experimental study on the chicken flexor tendons. Journal of Surgical Research 113, 195–200, doi:10.1016/S0022-4804(03)00204-X (2003).

53 Liang, J.-I. et al. Video-based Gait Analysis for Functional Evaluation of Healing Achilles Tendon in Rats. Annals of Biomedical Engineering 40, 2532–2540, doi:10.1007/s10439-012-0619-z (2012).

54 Nakajima, T. et al. Grafting of iPS cell-derived tenocytes promotes motor function recovery after Achilles tendon rupture. Nature Communications 12, 5012, doi:10.1038/s41467-021-25328-6 (2021).

55 Correa, S., Grosskopf, A. K., Klich, J. H., Hernandez, H. L. & Appel, E. A. Injectable liposome-based supramolecular hydrogels for the programmable release of multiple protein drugs. Matter 5, 1816–1838, doi:10.1016/j.matt.2022.03.001 (2022).

56 Meany, E. L. et al. Injectable Polymer-Nanoparticle Hydrogel for the Sustained Intravitreal Delivery of Bimatoprost. Advanced Therapeutics 6, 2200207, doi:10.1002/adtp.202200207 (2023).

57 Ou, B. S. et al. Broad and Durable Humoral Responses Following Single Hydrogel Immunization of SARS-CoV-2 Subunit Vaccine. Advanced Healthcare Materials 12, 2301495, doi:10.1002/adhm.202301495 (2023).

58 Han, G.-D. et al. Potent anti-adhesion agent using a drug-eluting visible-light curable hyaluronic acid derivative. Journal of Industrial and Engineering Chemistry 70, 204–210, doi:10.1016/j.jiec.2018.10.017 (2019).

59 Kulick, M. I., Brazlow, R., Smith, S. & Hentz, V. R. Injectable Ibuprofen: Preliminary Evaluation of Its Ability to Decrease Peritendinous Adhesions. Annals of Plastic Surgery 13, 459 (1984).

60 Liu, S. et al. Prevention of Peritendinous Adhesions with Electrospun Ibuprofen-Loaded Poly(l-Lactic Acid)-Polyethylene Glycol Fibrous Membranes. Tissue Engineering Part A 19, 529–537, doi:10.1089/ten.tea.2012.0208 (2013).

61 Branford, O. A., Klass, B. R., Grobbelaar, A. O. & Rolfe, K. J. The growth factors involved in flexor tendon repair and adhesion formation. Journal of Hand Surgery (European Volume) 39, 60–70, doi:10.1177/1753193413509231 (2014).

62 Mao, W. F. et al. Modulation of digital flexor tendon healing by vascular endothelial growth factor gene transfection in a chicken model. Gene Therapy 24, 234–240, doi:10.1038/gt.2017.12 (2017).

63 Freedman, B. R. et al. Enhanced tendon healing by a tough hydrogel with an adhesive side and high drug-loading capacity. Nature Biomedical Engineering 6, 1167–1179, doi:10.1038/s41551-021-00810-0 (2022).

64 Meany, E. L. et al. Generation of an inflammatory niche in an injectable hydrogel depot through recruitment of key immune cells improves efficacy of mRNA vaccines. bioRxiv, 2024.2007.2005.602305, doi:10.1101/2024.07.05.602305 (2024).

