## Supplementary Material for "Preventing peritendinous adhesions using lubricious supramolecular hydrogels"

### 1. Supplementary Methods

*Materials:* Hypromellose (HPMC, meets USP testing specifications), N-methyl-2-pyrrolidone (NMP), 1-dodecylisocynate, N,N-diisopropylethylamine (DIPEA), acetone, monomethoxy-PEG (5 kDa), diazobicylcoundecene (DBU), acetic acid, diethyl ether, hexanes, dimethyl sulfoxide (DMSO), acetonitrile were purchased from Sigma-Aldrich and used as received. Dichloromethane (DCM) was purchased from Sigma-Aldrich and further dried via cryo-distillation. Lactide was purchased from Sigma-Aldrich and recrystallized from ethyl acetate (dried over sodium sulfate) three times.

*HPMC-$C_{12}$ synthesis:* HPMC−$C_{12}$ was prepared according to previously reported procedures.[63] HPMC (1.0 g) (SEC MALS: Mw (Đ) = 372.4 kDa (1.43), method previously reported) was dissolved in NMP (40 mL) at room temperature with stirring. Once the polymer had completely dissolved, the reaction was brought to 50 °C and a solution of 1-dodecylisocynate (0.5 mmol) in NMP (5 mL) was added dropwise, followed by DIPEA (catalyst, 125 µL). The reaction was maintained at 50 °C for 30 minutes, then heat was shut off and mixture was left stirring overnight at room temp. The solution was then precipitated from acetone and HPMC-$C_{12}$ was purified by dialysis against MilliQ water for 3-4 days (MWCO 3.5 kDa) and lyophilized, yielding HPMC-$C_{12}$

as a white amorphous powder. The polymer was dissolved at 20 mg $mL^{-1}$ in sterile PBS, pH 7.4, prior to use in hydrogels. For NIR and rhodamine tagged HPMC-C12, synthesis followed reported protocol in main text methods.

*PEG-PLA synthesis:* PEG-PLA was prepared and analyzed as previously reported.[63] Recrystallized lactide (10 g) was fully dissolved in cryo-distilled DCM (45 mL) under $N_{2\,(g)}$ with mild heating. Methoxy poly(ethylene glycol) (5 kDa; 2.5 g) was heated to 100 °C under vacuum for 1-2 h, allowed to cool under $N_2$, and then dissolved in cryodistilled DCM (5 mL). Once dissolved, the full PEG solution was added to the lactide solution under $N_2$ and mixed with hand swirling. A solution of DBU (150 µL cryodistilled DBU per 1 mL cryodistilled DCM) was prepared and 500 µL added to the lactide-PEG solution under $N_2$. The reaction was swirled by hand and allowed to react for 8 min before quenching with acetic acid (~2 drops in 500 µL acetone). The PEG-PLA copolymer was precipitated from excess 50:50 mixture ethyl ether and hexanes, collected, and dried under vacuum to yield a white amorphous powder. DMF GPC: Mw (Đ) = 24.5 kDa (1.13), method previously reported.

*PEG-PLA nanoparticle (NP) preparation*: NPs were prepared and analyzed as previously reported (*34*). Briefly, a solution (1 mL) of PEG-PLA in 25:75 DMSO:acetonitrile (50 mg $mL^{-1}$) was added dropwise to water (10 mL) at a stir rate of 600 rpm. NPs were purified by ultracentrifugation over a filter (MWCO 10 kDa; Millipore Amicon Ultra-15) followed by resuspension in PBS to a final concentration of 200 mg $ml^{-1}$. NP size and dispersity were characterized by DLS (Wyatt DynaPro PlateReader-II; average diameter = 34.1 nm, PDI = 0.05).

*PNP hydrogel formulation:* HPMC-$C_{12}$ was dissolved at 6 wt% in PBS and loaded into a 1 mL luer-lock syringe. A 20 wt% solution of PEG-PLA NPs in PBS was loaded into a second 1 mL syringe. The two syringes were connected with a female-female luer lock elbow, with care to avoid air at the interface of the HPMC-C12 and nanoparticle solution, and gently mixed until a homogenous PNP hydrogel was formed. Hydrogels were formulated with final concentrations of 1 wt% HPMC-$C_{12}$ and 10 wt% NPs, denoted PNP-1-10.

*Estimation of shear strain in peritendinous space:* The minimum shear strain was estimated by approximating the tendon and synovial sheath as parallel plates sliding past one another. The length of flexor tendon excursion was divided by the maximum gap between tendon and sheath. Values for tendon excursion and gap between tendon and sheath were estimated based on published studies.[25-27]

### 2. Supplementary Data

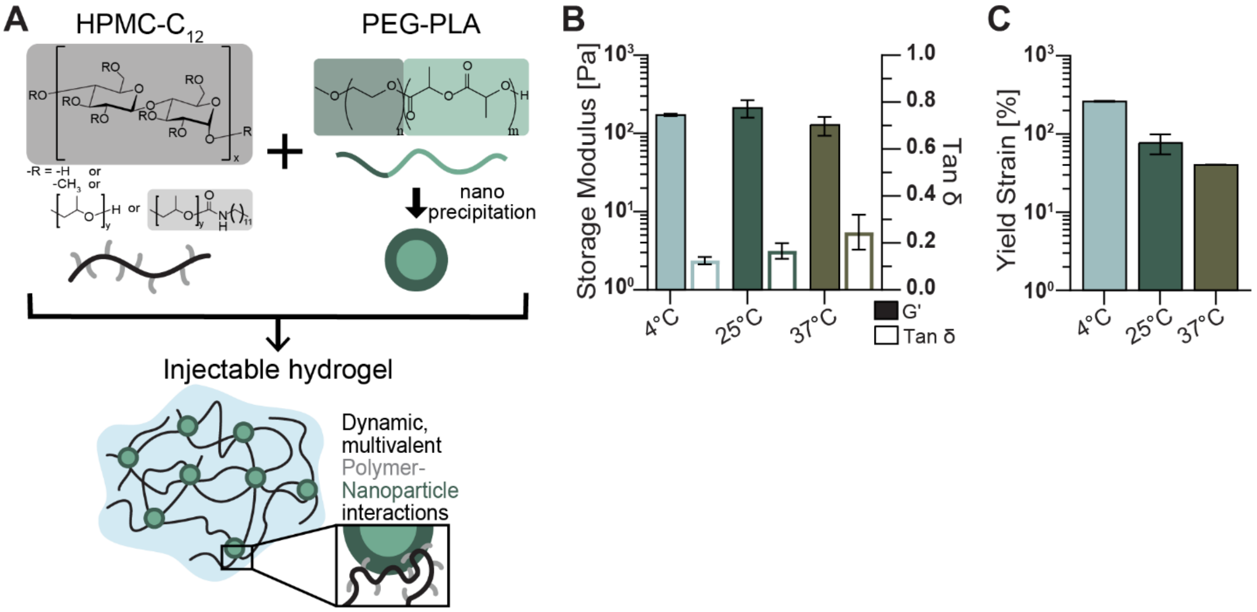


**Figure S1.** PNP hydrogel mechanical characterization. A) PNP hydrogels comprise HPMC-$C_{12}$ and PEG-PLA NPs. Multivalent hydrophobic interactions between dodecyl groups and nanoparticles create dynamic crosslinks. B) G′ and tan δ at 40 rad·$s^{-1}$ and C) yield strain (strain where G′ < 0.85 * max(G′)) for PNP at 4, 25, 37 °C. Data shown as mean ± SD, n=3.

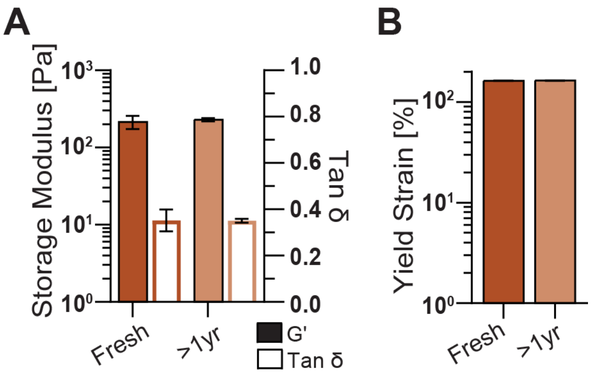


**Figure S2.** PTw mechanical properties following prolonged storage. A) G′ and tan δ at 40 rad·$s^{-1}$ and B) yield strain (strain where G′ < 0.85 * max(G′)) for PTw at 25 °C immediately after synthesis and after more than 1 year of storage at 4 °C. Data shown as mean ± SD, n=3.

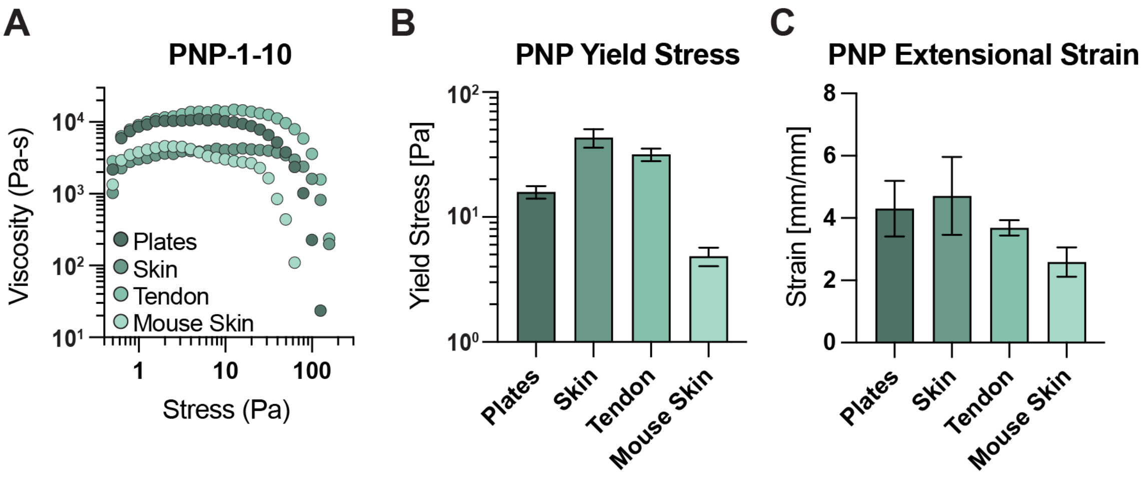


**Figure S3.** Tissue adhesiveness of PNP-1-10 hydrogel. A) Yielding behavior of PNP hydrogel in a serrated parallel plate geometry and on various tissue substrates in a stress-ramp experiment. B) Yield stress values of the PNP hydrogel defined at a 15% decrease from the peak viscosity. C) Extensional strain at break of PNP hydrogels before cohesive failure with various substrates. Data shown as mean ± SD, n=3.

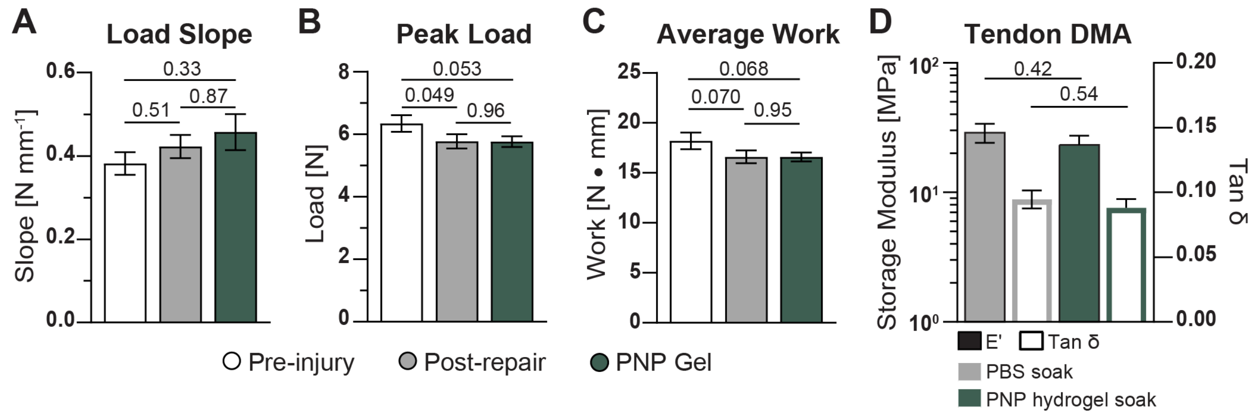


**Figure S4.** PNP hydrogel impact on human cadaver digits. A) Extracted load slope, B) peak load, and C) average work across digits. Data presented as mean ± SEM, n=5 - 8 across twelve unique digits from four arms and two patients. Statistical values shown are *p* values obtained from GLM fitting and Tukey HSD multiple comparison test in JMP (blocking by digit). D) extracted values of E′ and tan δ at 40 rad·s$^{-1}$ showing relative stiffness and elasticity of tendons is not impacted by PNP hydrogel material compared with PBS. Data shown as mean ± SEM, n=3 and statistical values are *p* values obtained from unpaired, two-tailed t-tests performed in GraphPad Prism.

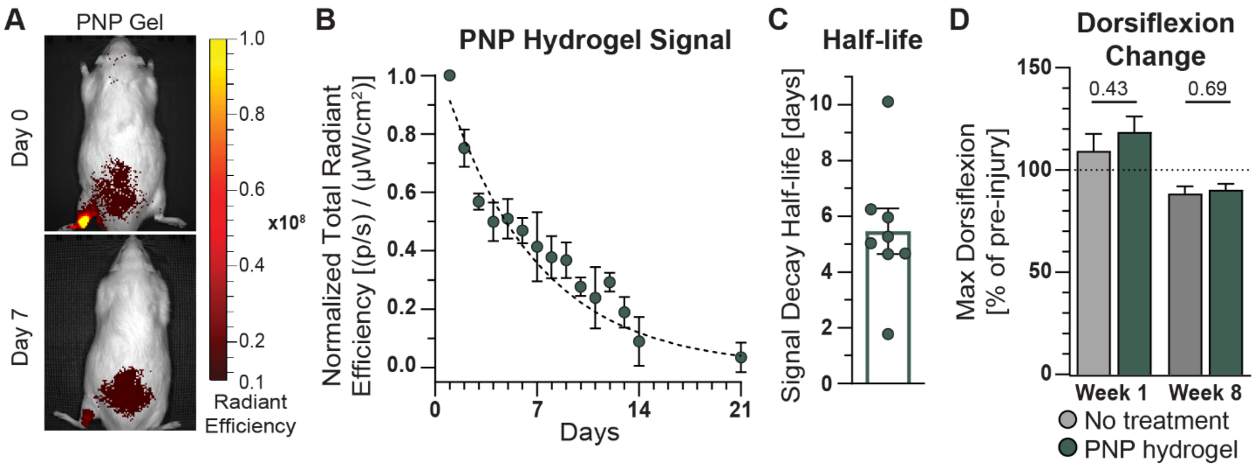


**Figure S5.** PNP hydrogel local retention and efficacy in rat Achilles tendon injury. A) Representative images of rats with PNP hydrogel on days zero and seven following surgery. B) Normalized fluorescent signal plotted over time along with C) extracted half-lives. D) Percent change in the maximal dorsiflexion angle for rats at weeks one and eight compared to the individual rat's pre-injury max dorsiflexion angle. Data presented as mean ± SEM, n=8. Statistical values are *p* values obtained from unpaired, two-tailed t-tests performed in GraphPad Prism.

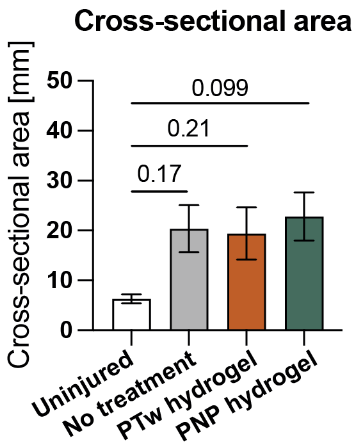


**Figure S6.** Cross-sectional area of harvested tendons from all treatment groups. Measured cross-sectional areas for uninjured and injured tendons. Data presented as mean ± SEM, n=3 for uninjured group, else n=5. Statistical values are *p* values obtained from GLM fitting and Dunnett's test in JMP with comparison to uninjured mean as control.

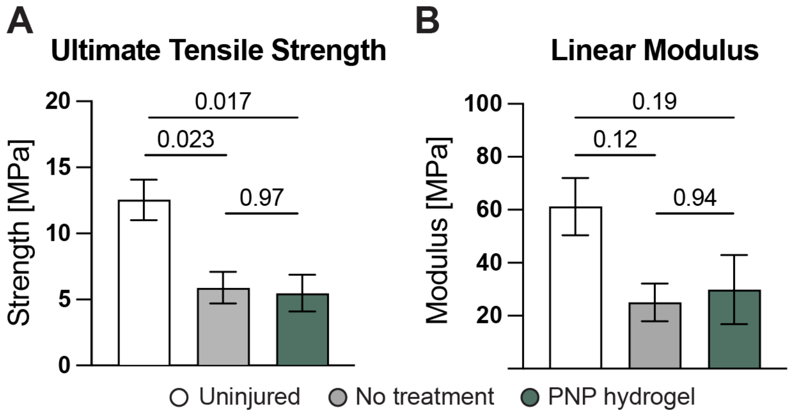


**Figure S7.** PNP impact on tensile strength of healing rat tendons. A) Ultimate tensile strength and B) linear modulus of tendons either uninjured, injured with no treatment, or injured with PNP hydrogel treatment. Data presented as mean ± SEM, n=3 for uninjured group, else n=5 and statistical *p* values obtained from GLM fitting and Tukey HSD multiple comparison test in JMP.

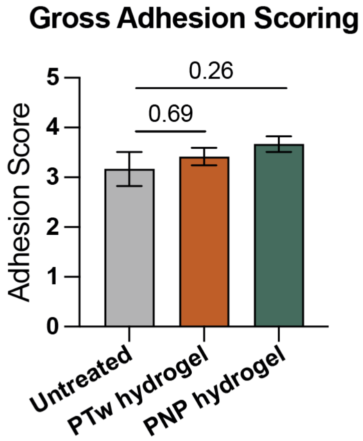


**Figure S8.** Double-blinded clinical scoring of gross adhesions eight weeks following Achilles tendon injury. Data presented as mean ± SEM, n=8 tendons, three independent scorers. Statistical values are *p* values obtained from GLM fitting and Dunnett's test in JMP with comparison to uninjured mean as control.

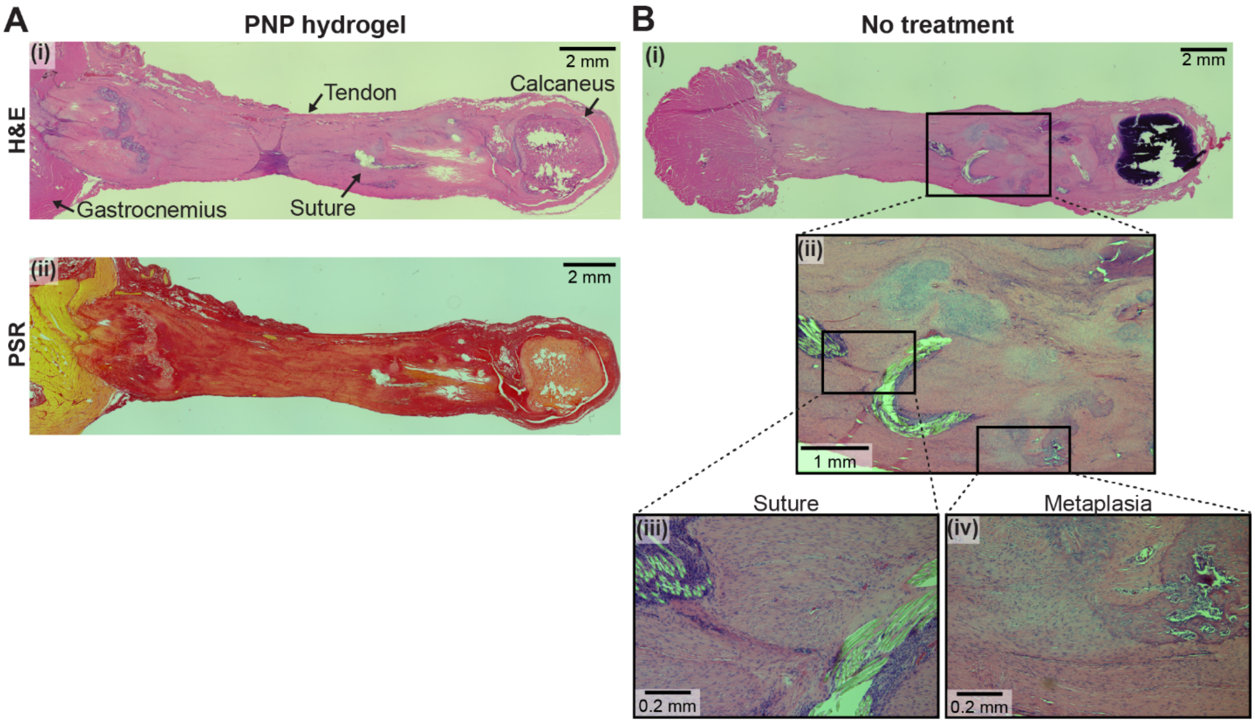


**Figure S9.** Histology of rat Achilles tendons at week eight. A) Representative images of H&E and PSR staining for PNP hydrogel treated tendon showing similar healing to no treatment and PTw treated tendons. B) Representative no treatment H&E stained tendon at (i) 2.5x with (ii) selected region and 10x zoomed images showing (iii) the local inflammatory response around suture material and (iv) chondroid (lighter pink region) and osseus metaplasia (dark region).

**Table S1.** Rat dorsiflexion angle metrics.

Dorsiflexion Metrics

| *Pre-Injury* | All Animals | No Treatment | PTw hydrogel |
|---|---|---|---|
| Mean | 49.2 | 55.2 | 45.3 |
| Median | 50.5 | 55.4 | 43.8 |
| IQR | 14.3 | 10.2 | 12.8 |

| *Week 8* | | No Treatment | PTw hydrogel |
|---|---|---|---|
| Mean | | 48.3 | 51.3 |
| Median | | 47.9 | 50.6 |
| IQR | | 7.5 | 4.1 |

**Table S2.** *p* values from ordinary one-way ANOVA with Dunnett's test and comparison to uninjured mean as control, run in GraphPad Prism, comparing rat dorsiflexion angles over time within treatment groups.

Dorsiflexion Angles

| No Treatment | | | Adjusted *p* value |
|---|---|---|---|
| Pre-injury | vs. | Week 1 | 0.46 |
| Pre-injury | vs. | Week 8 | 0.086 |

| PTw hydrogel | | | Adjusted *p* value |
|---|---|---|---|
| Pre-injury | vs. | Week 1 | ***0.0004*** |
| Pre-injury | vs. | Week 8 | 0.14 |

| PNP hydrogel | | | Adjusted *p* value |
|---|---|---|---|
| Pre-injury | vs. | Week 1 | 0.10 |
| Pre-injury | vs. | Week 8 | 0.32 |

**Table S3.** Adhesion scoring criteria.

Adhesion score criteria

| Score | Criteria |
|---|---|
| 1 | No or nearly no adhesion |
| 2 | Adhesion area can be separated by blunt dissection alone |
| 3 | Adhesion area is less than 50%, which requires sharp dissection for separation |
| 4 | Adhesion area is more than 50%, which requires sharp dissection for separation |
| 5 | Adhesion area of 97.5% or more, which requires sharp dissection for separation |
